# Integrative Analysis of Drug-Gene Expression Signatures in Human Pluripotent Stem Cells Identifies Novel Drug Candidates for ALS and Monogenic Diseases

**DOI:** 10.1101/2025.02.04.636445

**Authors:** Florine Roussange, Jacqueline Gide, Johana Tournois, Michel Cailleret, Anne Boland, Christophe Battail, Jean-François Deleuze, Hélène Polvèche, Didier Auboeuf, Knut Brockmann, Edor Kabashi, Anca Marian, Lina El Kassar, Sophie Blondel, François Salachas, Gaëlle Bruneteau, the PULSE study group, Marc Peschanski, Cécile Martinat, Sandrine Baghdoyan

**Author notes:** Jacqueline Gide died in April 2022.

## Abstract

The classical paradigm of drug screening often faces significant limitations due to the challenges associated with identifying molecular or cellular read-outs that are relevant to specific diseases. This issue is particularly pronounced for the thousands of diseases of genetic origin, given the abundance of databases listing disease-associated changes in gene expression and various transcripts, indicating that potential read-outs may be concealed within these resources. To remedy this, an alternative approach was tested: compounds were evaluated for their effects on gene expression and alternative splicing in a healthy cell model, and the resulting data were matched to molecular signatures of diseases. A subset of 50 FDA-approved drugs was tested on mesenchymal stem cells derived from a human pluripotent stem cell line. Over half of the compounds altered gene expression, many affecting pathways linked to monogenic diseases. One hit, increased SQSTM1 expression induced by prazosin, was further validated in ALS models caused by SQSTM1 haploinsufficiency, including patient-derived fibroblasts, SQSTM1-depleted hiPSC-derived motor neurons, and a zebrafish model. Extending this paradigm could involve testing diverse cell types and larger drug libraries.

**One Sentence Summary:** A novel drug screening approach for monogenic diseases integrating human pluripotent stem cell derivatives and RNA sequencing to profile gene expression.

## INTRODUCTION

Drug discovery has long been the cornerstone of medical progress, traditionally focused on testing a wide range of pharmacological compounds against a predefined target, or “read-out,” linked to a specific disease. While this paradigm has achieved notable successes, it faces significant limitations, mainly due to its single target approach to addressing a specific pathology (1, 2). As a result, the scope of discovery is largely confined to the interaction between a drug and a target, often under highly controlled conditions, which limits consideration of the wider biological context. Indeed, the diversity in classical drug screening depends largely on the pharmacological compounds tested and their application parameters (e.g., dosage, timing). This fails to fully exploit the rich biological variability within cellular systems and molecular mechanisms, critical factors especially for diseases driven by complex changes in gene expression and regulatory networks. Beyond these issues, classical drug screening faces difficulties in effectively exploiting the wealth of data provided by modern genomic and transcriptomic databases. Advances in high-throughput sequencing technologies and computational resources have generated vast datasets cataloguing gene expression profiles and splicing variations in many diseases (3, 4). These data provide valuable information that could significantly improve drug discovery if properly integrated into screening strategies. Initiatives such as the Broad Institute’s Connectivity Map have begun to generate comprehensive resources for analyzing gene expression changes in various tumor cell types in response to specific compounds (5, 6). This widely used database has been instrumental in identifying potential agents that target key cancer-related pathways (7–10).

Such an approach offers significant potential for addressing rare genetic diseases, where a single gene mutation or a well-defined pathway is often at the root of the condition. By focusing on the cell types most pertinent to these specific pathologies, it becomes possible to target more relevant therapeutic mechanisms. The effectiveness of this strategy can be further amplified when combined with repositionable drugs -compounds that have already been approved by regulatory agencies for other diseases-speeding up the drug development process and increasing the likelihood of finding effective therapies (11). However, to take full advantage of this potential, it is necessary to change the paradigm of drug discovery. Rather than focusing solely on a specific disease, an agnostic approach to drug discovery could be adopted, whereby compounds are tested first and then the observed changes in gene expression and splicing are linked to relevant diseases.

In this study, we explored such an alternative approach to the classical drug screening paradigm by evaluating the potential of integrating human pluripotent stem cell derivatives with deep RNA sequencing to map gene expression signatures induced by 50 FDA-approved drugs. More than half of the compounds influenced gene expression, and many affected pathways linked to monogenic diseases. To assess the predictive value of this approach, we focused on prazosin, an antihypertensive drug identified as a modulator of SQSTM1 expression. SQSTM1 encodes the autophagy receptor SQSTM1/p62, mutations in which are linked to rare forms of amyotrophic lateral sclerosis (12, 13). Using patient-derived fibroblasts, SQSTM1-depleted hiPSC-derived motor neurons and an in vivo zebrafish model with sqstm1 knockdown, we demonstrated that prazosin treatment can enhance autophagy-related processes and improve motor neuron phenotypes, correlating with improved motor behavior in vivo.

## RESULTS

### Gene expression signature induced by 50 FDA-approved small molecules

A collection of 50 FDA-approved small molecules was selected based on the following criteria: (i) frequent prescription, (ii) known safety profiles, and (iii) diversity in chemical structure and biological activity (**Table S1**). To evaluate the transcriptional effects of these drugs, hES-derived mesodermal progenitor cells (MPCs) were utilized, as they have been previously established as suitable for drug screening applications (*14*) (**Fig. 1A**). Next-generation sequencing was performed on cells treated for 24 hours with a single concentration of 10 μM, a condition confirmed to be non-toxic (**Fig. S1**). The potential of these compounds was evaluated by analyzing their capacity to influence both transcript expression and alternative splicing. The percentage of alternative spliced values was estimated by systematic comparison of a mapping-first approach (FARLINE (*15*)) using a threshold of at least 10% change of the mean value. This analysis indicated that 74% of the drugs exerted a limited effect on RNA splicing, with only 10 to 50 events displaying a ΔPSI (Percentage Spliced In) value greater than 10% (**Fig. 1B** and **Table S2**). Only seven splicing events, out of 2,071 detected, exhibited ΔPSI values exceeding 50%. Three compounds - pentamidine, manumycin A, and rosuvastatin-emerged as having notable regulatory effects, each affecting more than 100 splicing events. Two of these drugs, namely manumycin A and pentamidine, had previously been identified as alternative splicing regulators with therapeutic potential for treating spliceopathies (*16, 17*). These regulatory effects were broadly distributed across various types of RNA splicing, with the exception of mutually exclusive exon splicing, which is observed in only 1% of cases (**Fig. 1C**). Although 88% of the compounds inducing splicing changes targeted genes linked to genetic diseases, none of the alternative splicing events was associated with a pathological process (**Fig. 1D**). Differentially expressed genes (DEGs) were selected on the basis of a false discovery rate (FDR) < 0.05 and Log2FoldChange > 0.4. Most of the compounds resulted in minimal transcriptional changes: 62% of the drugs affected the expression of fewer than 10 genes. Analysis of the DEGs revealed that approximatively 20% had previously been reported as mutated in a monogenic disease (**Fig. 1D**). Analysis by RT-qPCR on mesodermal progenitors confirmed that the expression of eight genes associated with pathogenic mutations was modulated by manumycin A, pentamidine, prazosin, and rosuvastatin (**Table 1 and Fig. 1E**). Interestingly, none of these compounds were robustly identified as modulators of these genes in the Broad Institute’s Connectivity Map database (**Fig. S2**). Since the results were derived from mesodermal derivatives, their relevance was further evaluated in the context of other cell types. For instance, several compounds were identified as modulators of genes associated with pathologies affecting ectodermal derivatives such as genodermatoses. RT-qPCR analyses, conducted on primary human keratinocytes, revealed variable outcomes depending on the genes, highlighting the critical role of the cell type of origin in determining pharmacological drug response (**Fig. S2**).

**Fig. 1.**
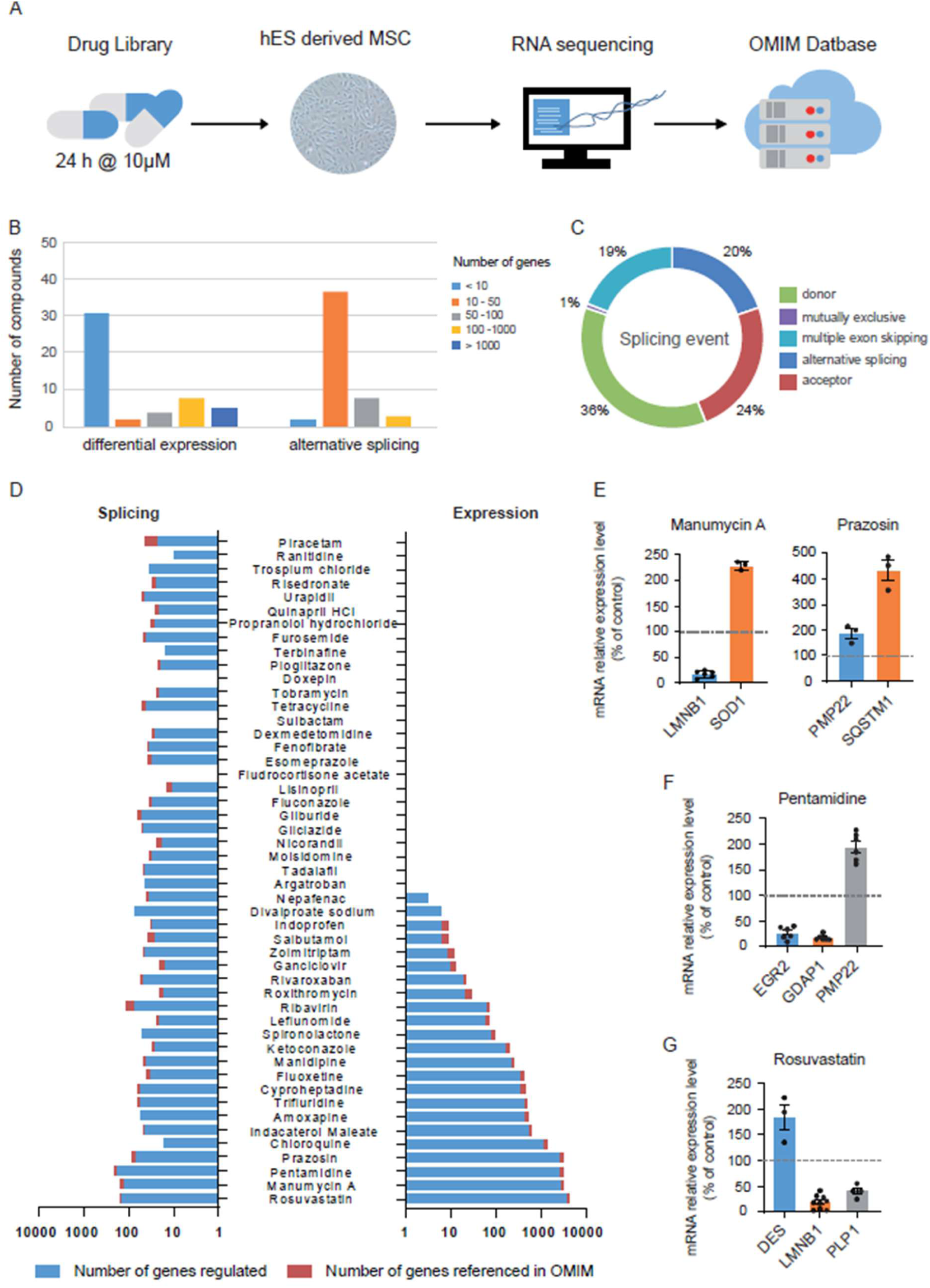
Molecular screening of repositioning drugs by RNA-seq identifies new therapeutic candidates for rare diseases. **(A)** Schematics illustrating the experimental paradigm. After 24h of treatment with drugs at 10µM, mRNAs from hES-derived MPCs were extracted and submitted to transcriptomic analysis to identify therapeutic regulation of genes involved in monogenic diseases, referenced in the OMIM database. **(B)** Graphs showing the number of drugs that regulate transcript expression (LEFT) or alternative splicing (RIGHT), categorized based on the number of events significantly altered compared to the control. **(C)** Distribution of the different types of alternative splicing events induced by drug treatments. **(D)** Graphs displaying the number of genes significantly regulated by each drug with log2 fold change >0.4 or ΔPSI > 10% (blue) as well as the proportion of genes referenced in OMIM (red). **(E-G)** Normalized expressions of *LMNB1, SOD1, PMP22, SQSTM1, EGR2, GDAP1, DES* and *PLP1* genes analysed by quantitative RT-PCR in hES-derived MPCs treated with different drugs at 10µM for 24h. Mean ± SD, n=3 experiments.

**Table 1.** Drugs regulating disease-causing genes.

| Gene | Disease |  | OMIM number | Transmission | Molecular Genetic | Incidence | Drug<br>Log2FC>1.45 | Regulation |
| --- | --- | --- | --- | --- | --- | --- | --- | --- |
| PLP1 | Pelizeus-Merzbacher Disease | PMD | 312080 | Recessive X linked | Duplication | 1/ 7663 | Rosuvastatin | Down |
| LMNB1 | Adult-onset Demyelinating Leukodystrophy | ADLD | 169500 | Autosomal dominant | Duplication | < 1 / 1 000 000 | Rosuvastatin | Down |
| SQSTM1 | Frontotemporal dementia / Amyotrophic Lateral Sclerosis 3 | FTDALS3 | 616437 | Autosomal dominant | Loss of function | 1-9 / 100 000 | Prazosin | UP |
| DES | Myofibrillar myopathy-1 | MFM1 | 601419 | Autosomal dominant | Loss of function | < 1 / 1 000 000 | Manumycin<br>Rosuvastatin | UP |

### Validation of drug-gene interactions in patient-derived cells with rare monogenic diseases

The ability of these compounds to modulate the expression of the targeted gene in a pathological context was next evaluated. However, due to the association of these genes with rare diseases, obtaining patient cells proved challenging. Fibroblasts were obtained only from patients carrying pathogenic mutations in the *PLP1* and *SQSTM1* genes (**Fig. 2A**).

**Fig. 2.**
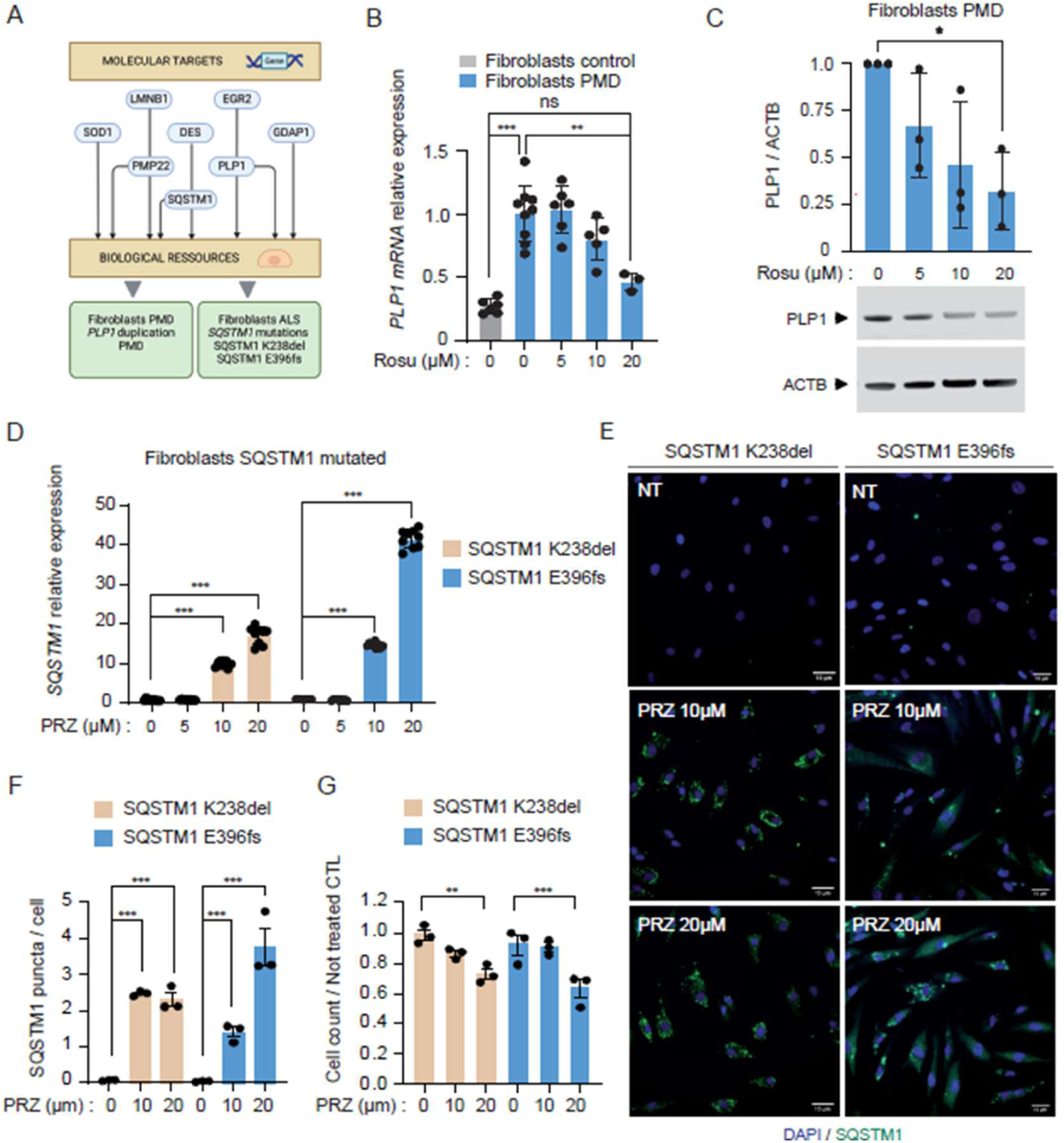
Rosuvastatin and prazosin promote therapeutic gene regulation of *PLP1* and *SQSTM1* transcripts in fibroblasts derived from PMD and ALS3 patients, respectively. **(A)** Schematic representation of the validation process for drug effects in fibroblasts derived from disease-affected patients. **(B)** Graphs showing *PLP1* transcript expression levels in PMD-fibroblasts, analyzed by RT-qPCR following treatment with varying doses of rosuvastatin, compared to non-affected fibroblasts. Data are presented as Mean ± SEM from three independent experiments (n=3); statistical analysis was performed using one-way ANOVA with Tukey’s post hoc multiple comparisons test. **(C)** Western blotting analysis of PLP1 and ACTB proteins in *PLP1* mutated fibroblasts treated with 5, 10 and 20µM of rosuvastatin. Data are shown as mean ± SEM of n=3 independent experiments, one way ANOVA with post hoc Dunnett’s multiple comparisons test. **(D)** Graphs of *SQSTM1* transcript expression in FTD/ALS fibroblasts from two patients (SQSTM1 K238del and SQSTM1 E396fs), analyzed by RT-qPCR after 24-hour treatment with various doses of prazosin. Data are represented as mean ± SEM from three independent experiments (n=3); one-way ANOVA with Dunnett’s post hoc multiple comparisons test was used for statistical analysis. **(E)** Representative immunocytochemistry images showing DAPI staining (blue) and SQSTM1 staining (green) in SQSTM1-mutated FTD/ALS fibroblasts (K238del and E396fs). Data are presented as Mean ± SEM from three independent experiments (n=3); statistical analysis was conducted using one-way ANOVA with Tukey’s post hoc multiple comparisons test. **(F-G)** Quantification of nuclei and SQSTM1 puncta per cell following 24-hour treatment with 10 or 20 µM prazosin, determined through automated microscope acquisition and analysis. Data are expressed as Mean ± SD from three independent experiments (n=3); statistical significance was evaluated using one-way ANOVA with Dunnett’s post hoc multiple comparisons test. *P < 0.05, **P < 0.01, ***P < 0.001.Mean ± SD of n=3 independent experiments, one way ANOVA with post hoc Dunnett’s multiple comparisons test. *P < 0.05, **P<0.01,***P < 0.001.

Mutations in the *PLP1* gene are linked to Pelizaeus-Merzbacher disease (PMD), a rare degenerative central nervous system disorder (*18, 19*). Fibroblasts with a *PLP1* gene duplication, known to cause toxic overexpression and oligodendrocyte dysfunction (**Fig. S3**) (*20*), were obtained. RT-qPCR analysis of *PLP1* transcript levels in PMD fibroblasts confirmed its overexpression compared to unaffected fibroblasts (**Fig. 2B**). Treatment of PMD fibroblasts with various doses of rosuvastatin for 24 hours demonstrated that the drug effectively reduced *PLP1* levels at both the RNA and protein levels in fibroblasts derived from PMD patients (**Fig. 2B-C**). Fibroblasts harboring SQSTM1 mutations (*K238del* and *E396fs*) associated to FTD/ALS3 were obtained from two patients (*12*). Due to the loss of function caused by these heterozygous mutations, prazosin treatment was expected to enhance the cellular levels of functional SQSTM1 produced by the non-mutated allele. An increase in SQSTM1 expression was confirmed after 24 hours of treatment with 10µM of prazosin using RT-qPCR analysis (**Fig. 2D**). This result was further validated at the protein level by quantifying the number of *SQSTM1* puncta detected by immunostaining, which also confirmed the absence of cytotoxic effects at that concentration (**Fig. 2E-G**). These results were further confirmed in muscle cells derived from biopsies of sporadic ALS patients (sporadic and SOD1 mutation), extending the effect of prazosin to other forms of the disease (**Fig S3**).

Since *SQSTM1* is known to play a key role in autophagy (*21*), a process central to ALS pathology, and prazosin has been shown to influence autophagy (*22, 23*), this compound was chosen for further investigation in the context of ALS.

### Prazosin regulates several genes involved in autophagy

Gene set enrichment analysis (GSEA) of the prazosin transcriptomic signature in hES-derived MPCs revealed an upregulation of genes associated with autophagosomes, lysosomes and endocytic vesicles (**Fig. 3A**). Among these, seven of these genes (*UBC, MAP1LC3B, HSPA1A, HSPA1B, HSPA5, HSPB8* and *HSPH1*) were identified as closely linked to SQSTM1 using STRING software (**Fig. 3B**). The increased expression of these genes was validated using RT-qPCR in both control and SQSTM1 mutant fibrobasts treated with different doses of prazosin (**Fig. 3B**). However, within the autophagic pathway, prazosin appears to selectively modulate specific genes. Notably, the expression of the autophagy receptor OPTN, involved in cargo recruitment to autophagosomes, remained unaffected by prazosin. This indicates that prazosin influences a subset of genes within the autophagy pathway.

**Fig. 3.**
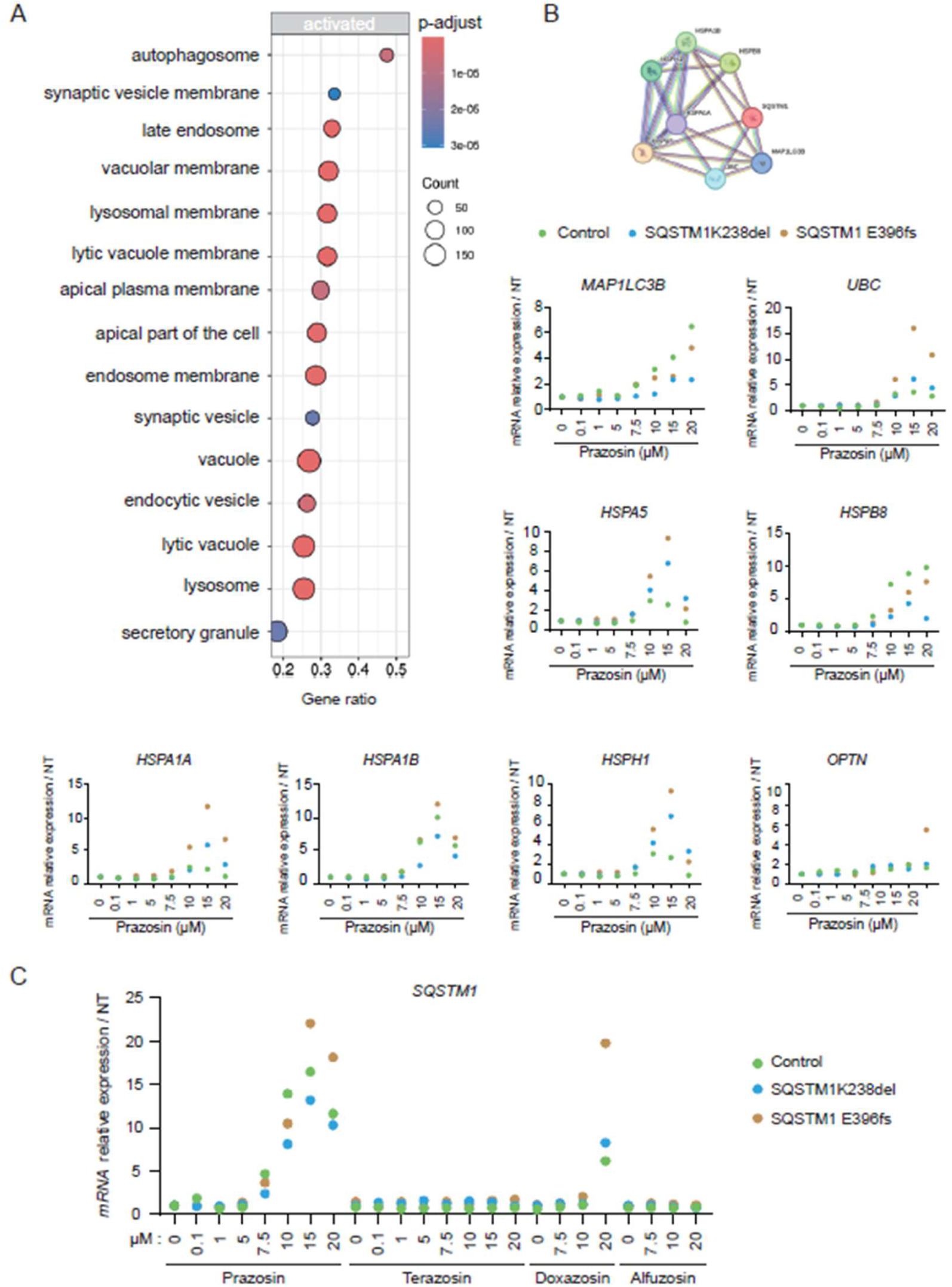
Prazosin regulates the expression of several genes involved in autophagy. **(A)** Gene set enrichment analysis (GSEA) of transcriptomic data from hES-derived MPCs treated with 10 µM prazosin for 24 hours reveals enrichment of genes associated with various organelles involved in the autophagic flux, including the autophagosome, late endosome, and lysosome. Data were analyzed using GSEA software to identify statistically significant enrichment in functionally related gene sets. **(B)** TRING network analysis of genes regulated by prazosin treatment in hES-derived MPCs (log2 fold change > 0.4) based on transcriptomic analysis after 24 hours of treatment with 10 µM prazosin. Quantitative real-time PCR analysis of *MAP1LC3B, UBC, HSPA5, HSPB8, HSPA1A, HSPA1B, HSPH1*, and *OPTN* gene expression was conducted in control fibroblasts and two SQSTM1-mutated fibroblast lines (#1 and #2) treated with varying doses of prazosin for 24 hours. **(C)** Quantitative real-time PCR analysis of *SQSTM1* gene expression in control fibroblasts and two *SQSTM1*-mutated fibroblast lines (#1 and #2) treated with different doses of prazosin, terazosin, doxazosin, and alfuzosin for 24 hours.

Prazosin is an antihypertensive agent that lowers blood pressure by selective alpha-1-adrenergic receptor antagonism. Recently, terazocin, another alpha-adrenergic receptor antagonist, has been shown to improve motor neuron phenotypes in several models of ALS by upregulating glycolysis and rescuing stress granule formation (*24*). We thus investigated whether terazosin, along with doxazosin and alfuzosin, two additional alpha-adrenergic receptor blockers, regulate *SQSTM1* expression. RT-qPCR analysis revealed that only prazosin could induce a pharmacological dose-dependent regulation of the *SQSTM1* gene in both control and *SQSTM1* mutant fibroblasts (**Fig. 3C**). These results suggest that prazosin induces specific cellular effects independently of the alpha-adrenergic receptors blockade it shares with other compounds of the same chemical family.

### Prazosin induces SQSTM1 expression and autophagy in hES-derived motor neurons

Since ALS primarily affects motor neurons (MNs) in the motor cortex, brainstem, and spinal cord, the impact of prazosin treatment on *SQSTM1* expression in human motor neurons was evaluated. Spinal MNs were generated from hESCs using a previously developed protocol that produces over 75% ISLET1-positive (ISL1+) MNs within 14 days (*25*). Treatment with 10 µM prazosin for 24 hours resulted in an upregulation of *SQSTM1* transcripts, with no observed cell toxicity. This result was further confirmed at the protein level by an increased number of *SQSTM1* puncta (**Fig. 4A-C**). The effect of prazosin treatment on the autophagosome marker Microtubule-Associated Protein 1 Light Chain 3 (LC3) was also examined. Western blot analysis demonstrated an increased conversion of LC3-I to its membrane-associated form LC3-II, indicative of autophagy induction, after 24 hours of treatment with 10 µM prazosin (**Fig. 4D**). These findings confirm that prazosin upregulates *SQSTM1* and induces autophagy in hES-derived spinal MNs.

**Fig. 4.**
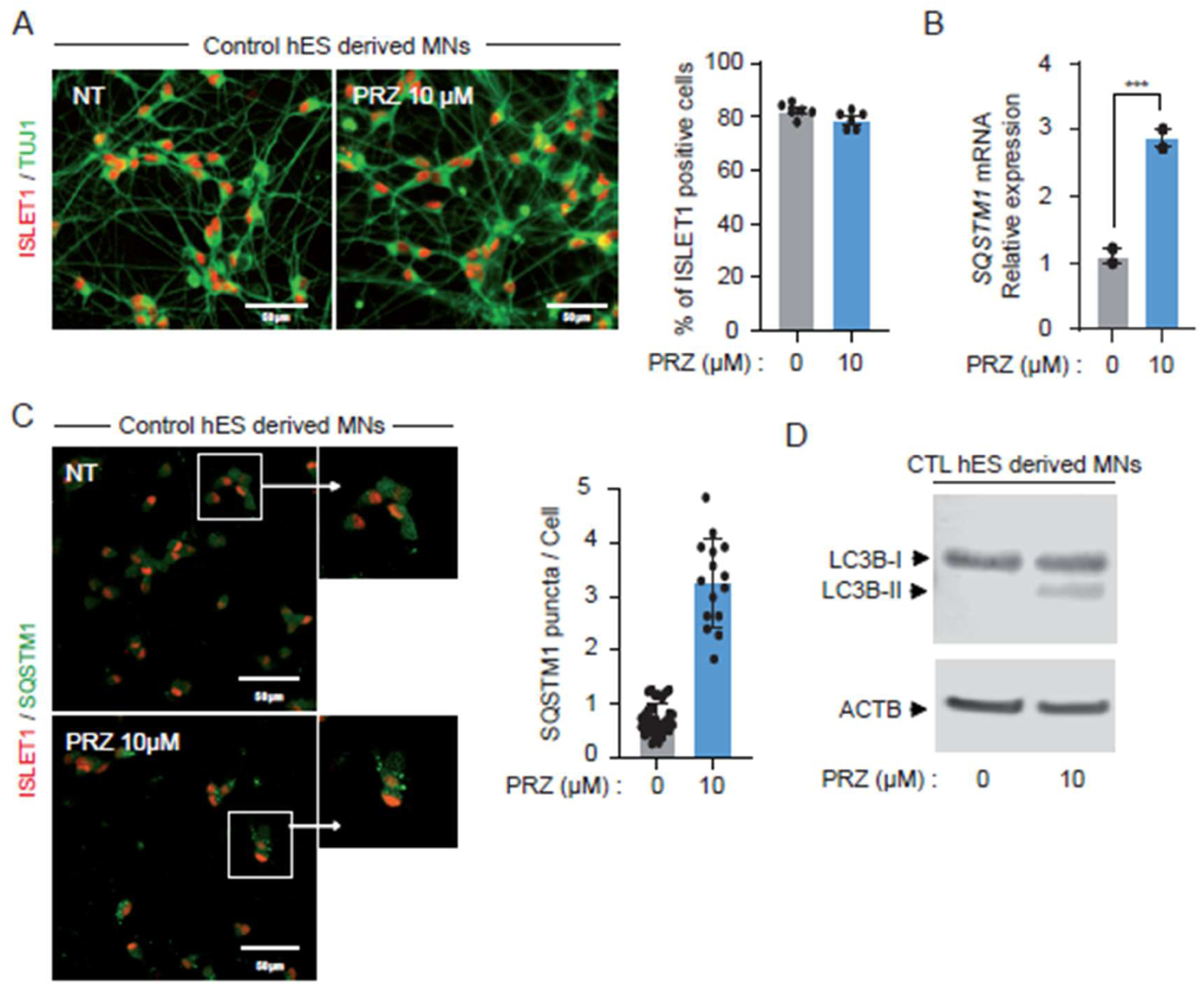
Prazosin regulates SQSTM1 expression and stimulates autophagy in hES derived motor neurons. **(A)** Representative immunocytochemistry images showing ISL1 (red) and TuJ1 (green) staining in control hES-derived motor neurons (MNs), either untreated (NT) or treated with 10 µM prazosin for 24 hours. The percentage of ISL1+ cells is presented as a graph based on data obtained through automated image acquisition and analysis. **(B)** Quantitative RT-PCR analysis of *SQSTM1* expression in hES-derived motor neurons (MNs) treated with 10 µM prazosin for 24 hours. Data are presented as mean ± SD from two independent experiments (n=2) and analyzed using Student’s *t*-test. **(C)** Representative immunocytochemistry images showing ISL1+ nuclei (red) and SQSTM1 puncta (green) staining in hES-derived MNs after 24-hour treatment with 10 µM prazosin. The number of SQSTM1 puncta per cell was quantified using automated microscope acquisition and analysis. **(D)** Western blot analysis of MAP1LC3B modification and ACTB expression in hES-derived MNs treated with 10 µM prazosin for 24 hours. The displayed image represents one of three independent experiments.

### Modeling SQSTM1 loss of function in hiPSCs derived motor neurons

We next sought to evaluate the therapeutic potential of prazosin in a more pathological context of SQSTM1 loss of function. Using CRISPR-Cas9 technology, different clones of hiPSC with SQSTM1 haploinsufficiency (SQSTM1 +/−) or knockout (SQSTM1 −/−) were generated (**Fig S4**). The pluripotency of edited clones was validated by immunostaining for the markers OCT3/4, NANOG, and SSEA3 and their genomic integrity was confirmed by karyotypic analysis (**Fig. S4**). Potential off-target activity of sgRNAs was assessed by analyzing the most likely off-target sites identified by CRISPOR and no genome-editing induced mutations were detected (**Fig. S5**). Two hiPSC clones of each genotype were differentiated into spinal MNs and SQSTM1 depletion was confirmed by western blot analysis (**Fig. 5A-C**). To assess the impact of SQSTM1 depletion on autophagy, SQSTM1 ^+/−^ and SQSTM1 ^−/−^ hiPSC-derived MNs were treated with the autophagic inducer torin-1. Western blot analysis revealed that the conversion of LC3-I to LC3-II in response to torin-1 treatment was significantly reduced in SQSTM1^+/−^ and SQSTM1^−/−^ MNs compared to control MNs (**Fig. 5D-E**). This result was further confirmed by analyzing the expression of LAMP2, another autophagy marker involved in lysosomal biogenesis and lysosomal fusion with autophagosomes. In control MNs, autophagy induction by Torin-1 resulted in reduced intensity of LAMP2 immunostaining whereas this effect was much less pronounced in SQSTM1+/− and SQSTM1−/− hiPSC-derived MNs (**Fig. 5F-G**).

**Fig. 5.**
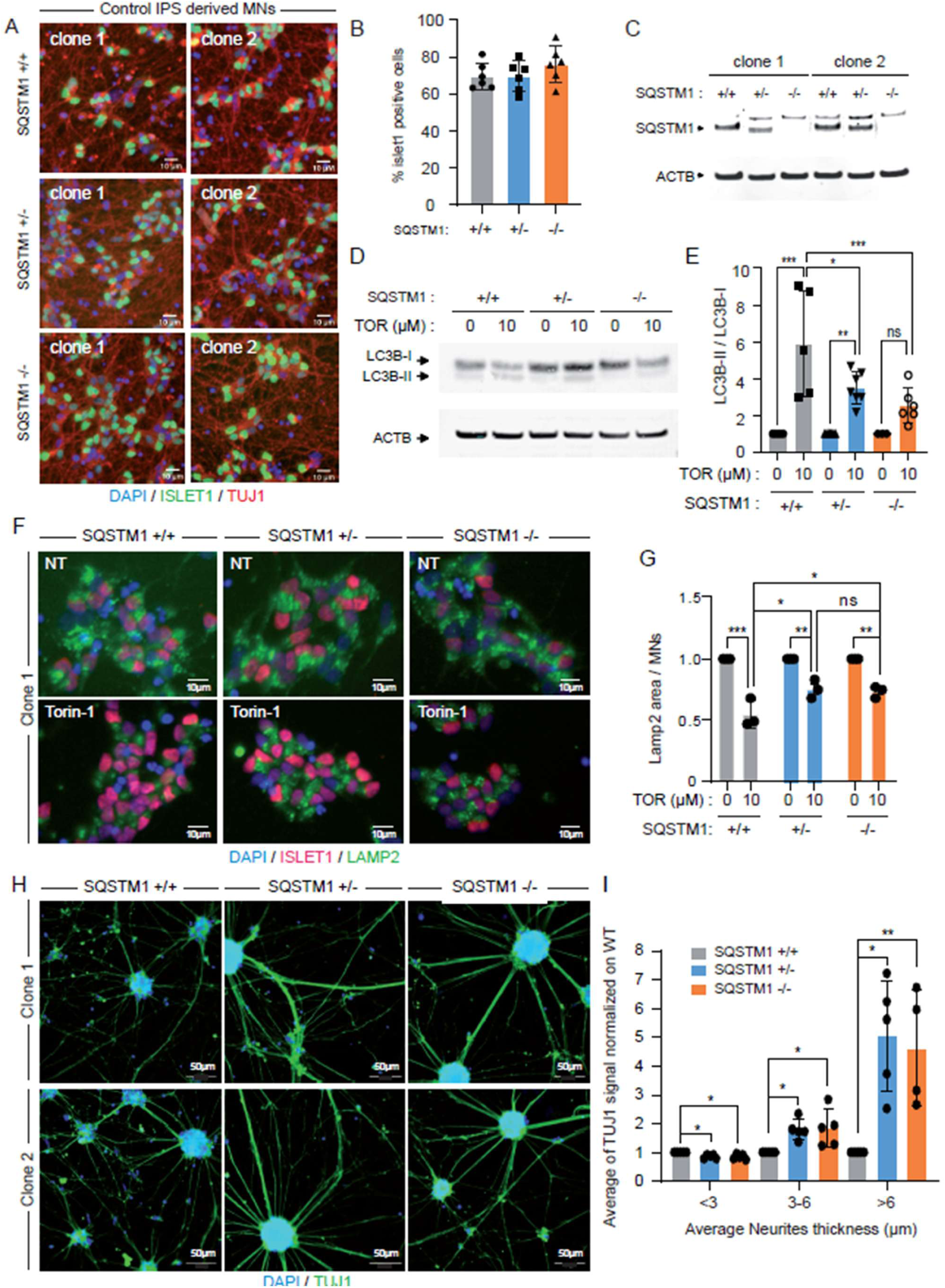
Motor neurons with heterozygous or homozygous SQSTM1 knockdown exhibit autophagy impairment and neuritic network perturbation. **(A)** Representative immunocytochemistry images showing DAPI (blue), ISL1 (green), and TuJ1 (red) staining in hiPSC-derived motor neurons (MNs) with *SQSTM1*+/+, *SQSTM1*+/−, and *SQSTM1*−/− genotypes after 14 days of differentiation. **(B)** Quantification of the percentage of ISL1+ MNs derived from control and *SQSTM1*-edited hiPSCs (two clones per genotype) using automated microscopic acquisition and analysis. **(C)** Western blot analysis of SQSTM1 and ACTB expression in MNs with *SQSTM1*+/+, *SQSTM1*+/−, and *SQSTM1*−/− genotypes derived from two edited hiPSC clones. **(D-E)** Representative images and quantification of MAP1LC3B modification analyzed by Western blot after 24-hour treatment with 10 µM Torin-1 in *SQSTM1*+/+, *SQSTM1*+/−, and *SQSTM1*−/− MNs derived from two individual edited hiPSC clones. Data are presented as mean ± SEM from more than three experiments (n > 3). **(F-G)** Representative immunocytochemistry images and quantification of DAPI (blue), ISL1 (red), and LAMP2 (green) staining in hiPSC-derived MNs with *SQSTM1*+/+, *SQSTM1*+/−, and *SQSTM1*−/− genotypes treated with 10 µM Torin-1 for 24 hours. Plots display the LAMP2 staining area normalized to ISL1+ MNs, presented as mean ± SD from more than three experiments (n > 3). Statistical significance was determined using two-way ANOVA with Šídák’s post hoc multiple comparisons test. **(H-I)** Representative immunocytochemistry images of DAPI (blue) and TuJ1 (green) staining in hiPSC-derived MNs with *SQSTM1*+/+, *SQSTM1*+/− and *SQSTM1*−/− genotypes after 24 days of differentiation. TuJ1 staining in MNs with different *SQSTM1* genotypes was quantified and normalized to *SQSTM1*+/+ MNs across three neurite thickness intervals. Data are presented as mean ± SD from more than three experiments (n > 3). Statistical significance was calculated using the Kruskal-Wallis test with Dunnett’s post hoc multiple comparisons test (*P < 0.05, **P < 0.01, ***P < 0.001).

Recent studies have reported defects in neuritic networks in ALS cellular models (*21, 26, 27*). Immunostaining for TUJI, a member of the beta-tubulin protein family, confirmed that neurites in SQSTM1+/− and SQSTM1−/− hiPSC-derived MNs tended to retract into large bundles compared to control hiPSC-derived MNs. By day 24 of differentiation, the width of these neurites is significantly increased in SQSTM1-depleted MNs (**Fig. 5H-I**). These observations suggest that decreased SQSTM1 levels in hiPSC-derived MNs impair autophagy and disrupt the structural integrity of the neurite network.

### Prazosin treatment rescues autophagic and neuritic phenotypes in SQSTM1^+/−^ hiPSC-derived motor neurons

The potential of prazosin to normalize the autophagic and neuritic phenotypes induced by SQSTM1 knockdown was then assessed. MNs derived from two SQSTM1+/− hiPSC clones were treated with 10µM prazosin for 24 hours. Immunostaining and Western blot analysis revealed that prazosin treatment enhanced the formation of SQSTM1 puncta and promoted the conversion of LC3-I to LC3-II in SQSTM1+/− hiPSC-derived MNs (**Fig. 6A-B**, **Fig. S6**). Prazosin treatment also resulted in a redistribution of lysosomes, as indicated by LAMP2 immunostaining, which shifted from the cell body to the neurites (**Fig. S6**). These findings suggest that prazosin may facilitate the translocation of lysosomes into neurites, thereby enhancing autophagy in axons. Consistently, prazosin treatment significantly reduced the formation of neurite bundles in SQSTM1+/− hiPSC-derived MNs without inducing toxicity, as indicated by the stable number of ISL1+ MNs compared to untreated conditions (**Fig. 6C-F**).

**Fig. 6.**
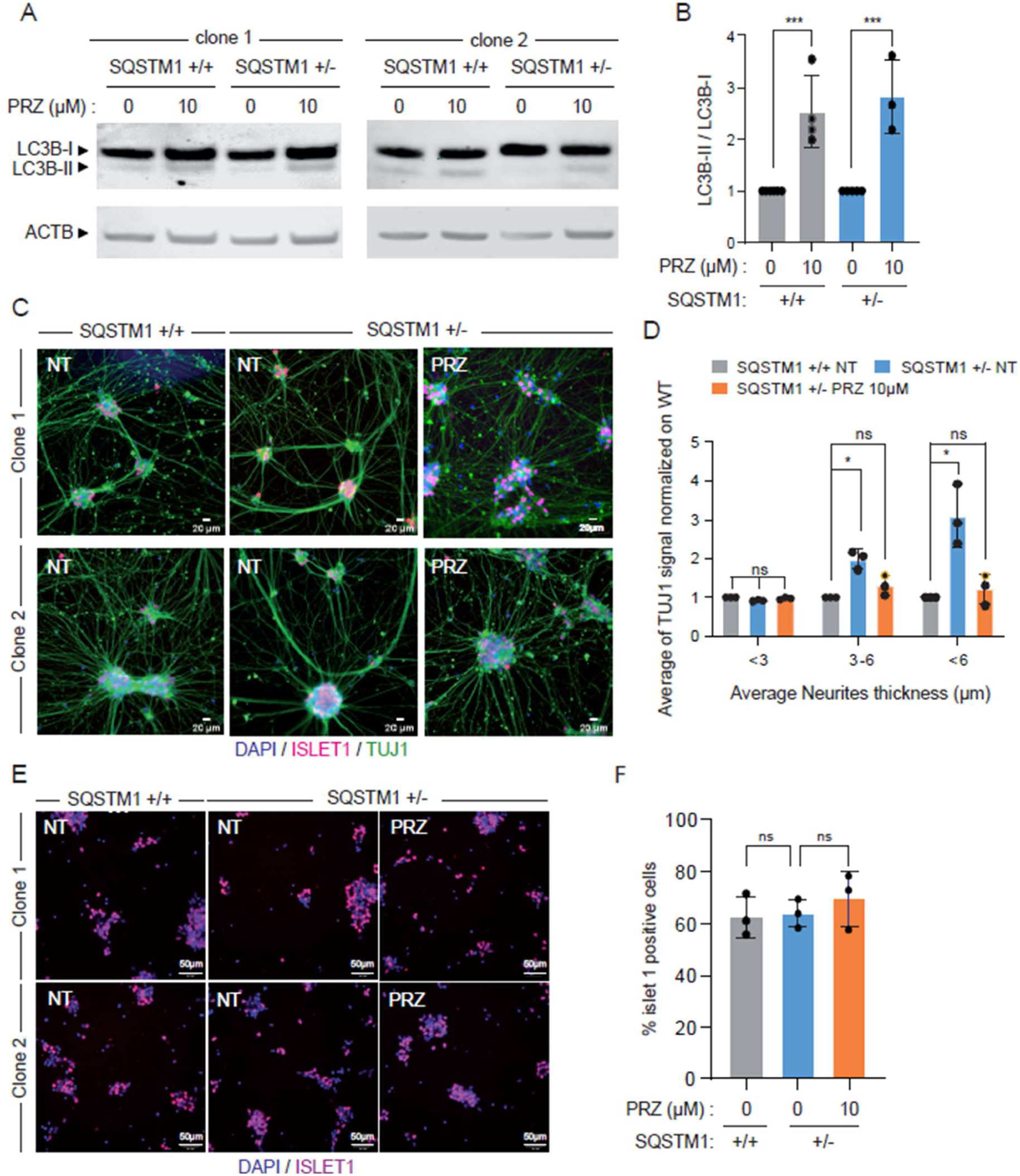
Prazosin promotes the autophagic flux in motor neurons with SQSTM1 haploinsufficiency. **(A-B)** Representative images and quantification of MAP1LC3B modification analyzed by Western blot in *SQSTM1*+/− motor neurons (MNs) derived from two individual edited hiPSC clones, treated with 10 µM prazosin for 24 hours. Data are presented as mean ± SEM from more than three experiments (n > 3). Statistical significance was determined using one-way ANOVA with Dunnett’s post hoc test (**P < 0.01). **(C-D)** Representative immunocytochemistry images and quantification of DAPI (blue) and TuJ1 (green) staining in MNs at 24 days of differentiation derived from *SQSTM1*+/+ and *SQSTM1*+/− hiPSCs, either untreated (NT) or treated with 10 µM prazosin for 10 days. TUJ1 staining in *SQSTM1*+/+ and *SQSTM1*+/− MNs was quantified and normalized to SQSTM1+/+ MNs across three neurite thickness intervals. Data are presented as mean ± SD from three independent experiments (n = 3). **(E-F)** Representative immunocytochemistry images and quantification of DAPI (blue) and ISL1+ (pink) staining in MNs derived from *SQSTM1*+/+ and *SQSTM1*+/− hiPSCs at 18 days of differentiation, either untreated (NT) or treated with 10 µM prazosin. Data are presented as mean ± SD from three independent experiments (n = 3). Statistical analysis was performed using the Kruskal-Wallis test with Dunnett’s post hoc multiple comparisons test (*P < 0.05, **P < 0.01, ***P < 0.001).

### Prazosin improves the swimming of zebrafish with sqstm1 knockdown

To assess the therapeutic potential of targeting SQSTM1 with prazosin in vivo, the drug was tested in a zebrafish model of ALS induced by SQSTM1 knockdown. As previously described (*28*), wild-type zebrafish embryos injected with an antisense morpholino oligonucleotide (AMO) targeting the sqstm1 ortholog exhibit shortened motor neuron axons and swimming deficits, which were quantified using the touch-evoked escape response (TEER) test (**Fig. 7A-B**). The absence of toxicity was first demonstrated after 4 days of treatment with prazosin in mismatch control embryos (**Fig. 7B**). Subsequently, sqstm1-AMO-injected zebrafish embryos were treated with prazosin, resulting in a significant improvement in swimming behavior during the TEER test compared to untreated littermates (**Fig. 7B**). This improvement was associated with an increase in sqstm1 protein expression (**Fig. 7C**).

**Fig. 7.**
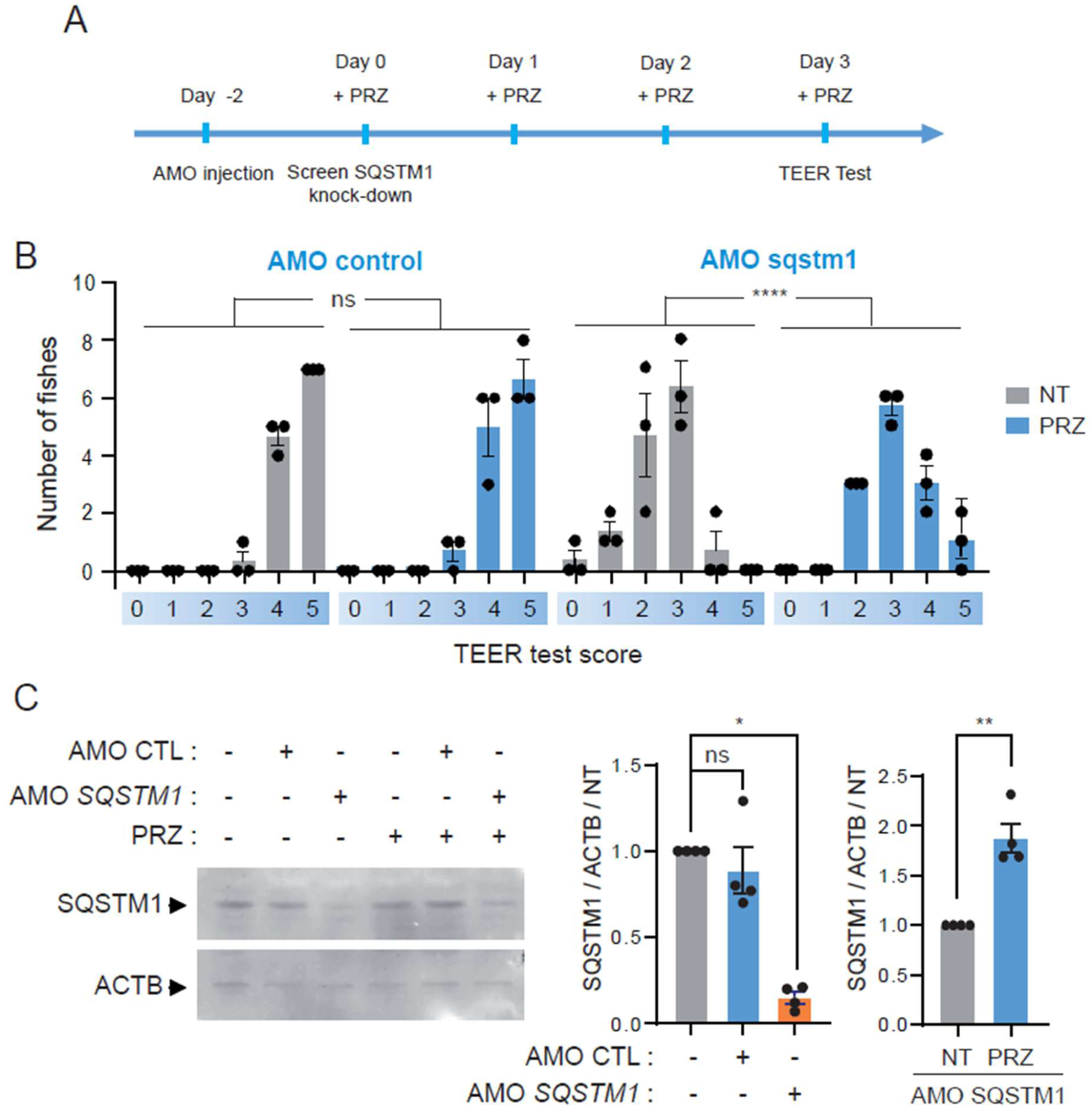
Prazosin alleviates the motor phenotype induced by *sqstm1* knockdown in zebrafish. **(A)** Schematic illustration of zebrafish treatment and functional assessment. **(B)** Quantification of swimming parameters in embryos injected with control or sqstm1-specific AMO, either untreated or treated with prazosin for 3 days, using a ViewPoint system. Data are presented as mean ± SEM from three independent experiments (n = 3). Statistical significance was determined using a Chi-square test for contingency data (****P < 0.0001). **(C)** Representative Western blot images and quantification of SQSTM1 and ACTB expression in embryos injected with control or sqstm1-specific AMO, either untreated or treated with prazosin for 3 days. Data are presented as mean ± SEM from three independent experiments (n = 3). Statistical analysis was performed using one-way ANOVA with Dunnett’s post hoc test and t-test (*P < 0.05, **P < 0.01).

## DISCUSSION

This study proposes an alternative to the traditional drug screening paradigm by combining human pluripotent stem cell derivatives with deep RNA sequencing to map gene expression changes induced by pharmacological compounds. This strategy highlights a systematic framework for advancing the development of personalized therapies, particularly applicable to monogenic diseases for which differentially expressed genes or an altered dosage of alternative transcripts have been identified. In the present study, we explored as proof of concept the potential therapeutic effect of prazosin, an antihypertensive drug, as a modulator of SQSTM1 expression, a gene encoding the autophagy receptor SQSTM1/p62, implicated in rare forms of ALS (*12, 13*). Using patient fibroblasts, SQSTM1-depleted hiPSC-derived motor neurons and a zebrafish sqstm1 knockdown model, prazosin has been shown to enhance autophagy processes and restore motor neuron phenotypes, including motor behavior in vivo.

Traditionally, hypothesis-driven approaches have been relied upon in drug repurposing, with cell-based assays used to explore overlaps between established pharmacological mechanisms and potential pathological pathways. The advent of disease-specific human pluripotent stem cells (hPSCs) has significantly advanced these methods (*29*). In the present study, a paradigm shift is introduced through the use of hPSCs in target-agnostic drug discovery approaches. Extending the scope of this research to a wider variety of hPSC-derived tissue types would be particularly beneficial in addressing the cell-type-dependent variability of pharmacological responses. Both our findings and the updated version of the Connectivity Map, which focuses on cancer cell lines, underline the importance of incorporating additional tissue types to better understand these responses (*6*). The ability of hPSCs to generate an unlimited supply of diverse cell types offers a powerful tool for exploring drug-specific gene expression profiles, potentially leading to the discovery of new therapeutic applications. Such a wide expansion would obviously benefit significantly from the integration of artificial intelligence (AI)-based methods to handle the vast amounts of data generated. Advances in AI are poised to address this challenge by enabling efficient data analysis and enhancing the scalability of these approaches (*30*). The combination of hPSC-derived cell diversity and AI-driven analytics promises to create a transformative platform for drug discovery, particularly for rare diseases for which the development of effective treatments has long been constrained by limited resources and inadequate biological models.

As a preliminary proof of concept for the paradigm shift in drug discovery we propose, we have demonstrated the potential to repurpose prazosin for a rare form of ALS. Currently, there is no effective cure for ALS, and the few treatments available slow disease progression down for only a few months. The heterogeneity of the disease, with numerous genetic mutations disrupting multiple molecular pathways, is one of the main challenges in developing a therapy for ALS. Over the past decades, research has highlighted the critical role of the autophagic pathway in ALS (*31*). Defects at different stages of autophagy are evident in experimental models of familial ALS, involving genetic mutations such as SOD1, SQSTM1, TDP-43, and OPTN, but probably also play an important role in the pathogenesis of sporadic ALS (*32*). As a result, targeting the autophagic pathway has emerged as a potential therapeutic approach for ameliorating ALS-related pathological conditions, and several compounds that positively modulate autophagy, such as trehalose, rapamycin, and colchicine, are already in clinical trials (*33–35*). Our results confirm the therapeutic potential of autophagy induction for ALS and identify prazosin as a promising new approach.

Recently, terazosin, a structural analog of prazosin, was shown to improve motor neuron phenotypes in several ALS models by activating the key glycolytic enzyme phosphoglycerate kinase 1 (*24*). Based on these results, a pilot clinical trial is currently underway to evaluate the potential of terazosin to reduce various biomarkers of neurodegeneration in ALS patients (*36*). Nevertheless, our results demonstrated that neither terazosin nor two other alpha-adrenergic receptor blockers, alfuzosin and doxazosin, inducedd a pharmacological increase in SQSTM1 expression. This suggests that prazosin regulates SQSTM1 gene expression independently of alpha-adrenergic receptors. Recent studies support this hypothesis by showing that prazosin can enter cells via endocytosis, leading to reorganization of the LAMP1-positive endolysosomal system (*37, 38*).

The effect of prazosin on autophagic processes, however, appears to extend beyond the regulation of SQSTM1 expression. Our deep RNA sequencing data indicate that prazosin modulates the expression of other autophagy regulators, including proteins known to interact with SQSTM1, such as UBC and MAP1LC3B. The molecular mechanisms by which prazosin regulates the expression of these different autophagy players remain speculative at this stage. It is possible that prazosin acts on a common mechanism that controls the expression or stability of these different genes. A potential candidate for this common mechanism could be TFEB, a master transcriptional regulator of genes involved in lysosome biogenesis and autophagy (*39*). However, our deep RNA sequencing data do not indicate that prazosin directly affects the expression of this gene. Further experiments are required to assess whether prazosin has an impact on post-transcriptional and post-translational regulation within this molecular pathway. Most importantly, these results suggest that prazosin broadly targets autophagy, paving the way for wider therapeutic applications.

Our study has certain limitations. Firstly, the precise mechanisms by which prazosin influences autophagic pathways remain unclear, requiring further experiments to elucidate its impact on transcriptional, post-transcriptional and post-translational regulation. Secondly, although our results suggest that prazosin has therapeutic potential for ALS, its efficacy and safety need to be validated by further preclinical and clinical studies, particularly in various genetic contexts and in sporadic ALS. Given that prazosin has already been tested in numerous clinical trials for repositioning in conditions such as traumatic stress (*40–43*) and Alzheimer’s disease (*44*), AI-driven analyses of data collected in these trials could provide valuable insights into its repositioning potential. Stratifying this data according to variables such as dosage and patient age could help uncover its true therapeutic potential. In addition, these approaches could also facilitate the selection of promising candidates for drug repurposing from the existing pharmacopoeia. Finally, although hPSC-derived models offer unique opportunities to assess drug effects, replicating the complexity of in vivo environments and predicting patient-specific responses remain significant challenges. The inclusion of more complex hPSC-derived models, such as organoids, could offer new perspectives to overcome these limitations and develop this paradigm into a robust and reliable platform for drug discovery, not only for ALS but also for other complex and rare diseases.

In conclusion, our systematic approach to drug-specific gene expression profiling in human pluripotent stem cell derivatives lays the foundation for the development of more effective and personalized therapeutic strategies.

## MATERIALS AND METHODS

### Chemicals

Small molecules listed in Supplemental Table 1 were purchased from Selleckchem. Prazosin (P7791), alfuzosin (A0232) and doxazosin (D9815) were purchased from Sigma-Aldrich®. Torin-1 (4247/10) and terazosin (1506/50) were purchased from Tocris.

### Origin and culture of primary cells from patients

All the primary cells obtained from patients were collected after informed consent had been obtained. All protocols were approved by the Institutional Review Boards at the institutions involved. Fibroblasts from a patient carrying a *PLP1* duplication and healthy fibroblasts were obtained from Dr. Knut Brockmann (Universitätsmedizin Göttingen, Germany). *PLP1* gene duplication was detected by SNP analysis. For that, high quality genomic DNA is obtained with QIAcube™ workstation using QIAamp® DNA Blood Mini Kit DNA Qiagen) from 5×106 cells. DNAg hybridation was achieved on Infinium Core-24v1-2 BeadChip (Illumina). Data were analyzed with GenomeStudio v2.0.5 software (Illumina). Fibroblasts carrying the K238 Del mutation in *SQSTM1* gene were provided by Dr Andrey Y. Abramov (UCL Institute of Neurology Queen Square, UK) (*45*). Fibroblasts carrying the mutation c.1184_1187dupTTGA (p.Glu396fs) in the *SQSTM1* gene were provided by Dr François Salachas (AP-HP, Paris ALS Centre, Hôpital de la Salpêtrière, Paris). Fibroblasts were cultured in DMEM (Invitrogen) supplemented with GlutaMAX (Invitrogen), 10% fetal bovine serum (Sigma-Aldrich) and 1% non-essential amino acids (Invitrogen) on 0.1% gelatin coated flasks. Primary myoblasts obtained from ALS patients were collected from the ongoing observational and prospective multicenter cohort PULSE (Study of Predictive Factors of Progression of Motor Neuron Disease). PULSE (Study of Predictive Factors of Progression of Motor Neuron Disease) is an ongoing observational and prospective multicenter cohort sponsored by the University Hospital of Lille, conducted in 16 ALS expert centers of the clinical research networks in France (FILSLAN, ACT4ALS-MND), approved by the CPP Nord Ouest-IV Ethics Committee (IDRCB number 2013-A00969-36) and registered on the ClinicalTrials.gov website (NCT02360891) (*46*). ALS-myoblasts were cultured in DMEM (Invitrogen) supplemented with GlutaMAX (Invitrogen), 20% fetal bovine serum (Sigma-Aldrich) and 1% Pyruvate de sodium (Gibco). Pooled primary human epidermal keratinocytes (HPEKp) were obtained from CELLnTEC and cultured in CnT-07 medium according to the manufacturer’s instructions.

### Differentiation, culture and treatment of hESC-derived MPCs

MPCs were differentiated from SA001 hESCs as previously described (*14*). This hESC line was used following the recommendation of the French Law of Bioethics and declared at the French Agency of Biomedicine (Number SSAB2133048S). MPCs were cultured on 0.1% gelatin-coated flasks and plates (SigmaAldrich) using Knockout Dulbecco’s Modified Eagle’s Medium (Invitrogen) supplemented with 20% fetal bovine serum (Eurobio, Les Ulis, France), 1mM Glutamax (Invitrogen), 1M non-essential amino acids (Invitrogen), and 0.1% β-mercaptoethanol (Invitrogen). For the RNA sequencing experiments, each compound was tested in triplicate at the dose of 10µM for 24 h. RNA were extracted from the cells using the RNeasy Micro Kit (Qiagen). The toxicity of the 50 compounds was assessed after 24h of treatment at 10 different doses using the CellTiter-Glo® Luminescent Cell Viability Assay (Promega).

### Generation of heterozygous and homozygous SQSTM1 knockout PSC lines

To generate *SQSTM1* depleted hiPSC clones, a control healthy hiPSC (1869) was engineered at the AAVS1 locus for doxycycline-inducible expression of SpCas9 protein using the homology-directed repair (HDR) donor plasmid pAAVS1-PDi-CRISPRn (Addgene #73500) (*47*). The generation and the use of this hiPSC line were approved by the french minister of health (2019-A02599-48). Briefly, hiPS cells were dissociated with StemPro Accutase Cell Dissociation Reagent (Gibco®), plated in 24-well plates at 25.000 cells per cm2 and treated with 50ng/mL doxycycline (Sigma-Aldrich®) to induce Cas9 expression. The next day, the cells were transfected with a mixed of 10pmol of sgRNAs targeting *SQSTM1* and tracrTNA using lipofectamine RNAiMax (Thermo Fisher Scientific®) according to the manufacturer’s protocol with appropriated guides targeting SQSTM1 and tracrRNA. sgRNAs targeting SQSTM1 sequence were determined by CRISPOR (http://crispor.tefor.net/) (Supplementary Table 3). To select edited clones, genomic DNA was extracted from transfected hiPS cells either with QIAmp DNA Micro and Mini Kit (Qiagen®) according to the manufacturer’s instructions. Gene editing was analyzed by Restriction fragment length polymorphism (RFLP) with the BtgZI enzyme (Supplementary Figure 3) and confirmed through Sanger sequencing. Genomic integrity was also verified by Multiplex fluorescence in situ hybridization (mFISH) karyotype analysis as described previously (*48*). To determine off-target activity of our gRNAs, we analyzed by PCR the most likely off-target sites predicted by CRISPOR (http://crispor.tefor.net/). No mutation induced by genome editing was observed (Supplementary Figure 4).

### Generation of spinal motoneurons from hiPSCs

The conversion of control and *SQSTM1* depleted hiPSC clones into spinal motoneurons was performed as previously described (*25*). Briefy, hiPSC were dissociated enzymatically using Stem Pro Accutase (ThermoFisher®) and plated in 25 cm2 fasks (Dutscher®) at 2×10^6^ cells per fask in an induced motoneuronal medium supplemented with cytokines every 2 days. After 10 days of differentiation, embryoid bodies were dissociated. Between days 10 and 14, MN progenitors are converted into MNs, and motoneuron phenotype was assessed by immunolabeling for ISLET1 (ISL1).

### RNA sequencing library preparation, sequencing

Sequencing libraries were prepared using the llumina TruSeq Stranded mRNA Sample Prep Kit (Illumina, San Diego, CA). A 2 × 101 bp paired-end sequencing was performed on the HiSeq2000 instrument, using half a lane per sample, to produce on average 80 million read pairs per sample (160 million sequences) with an average insert length of 130 bp. The samples were sequenced at this depth to provide sufficient coverage for gene expression and alternative splicing analyses. Trimmomatic (*49*), Tophat2 (*50*), Picard suite (http://www.broadinstittute.github.io/picard), RNA-SeQC (*51*) and in-house metrics were used to evaluate data quality. The quality control of the sequencing data was evaluated using FastQC. Reads were aligned using TopHat2 (v2.0.853). TopHat2 was run with the assistance of gene annotations (Illumina’s iGenomes based on EnsEMBL r70), which means that the alignment was performed in three steps: transcriptome mapping, genome mapping, and spliced mapping.

### Bioinformatic analysis of splicing events and differential genes expression

RNA-seq data from three biological replicates for each condition were analysed using FaRLine (FasterDB RNAseq Pipeline) in order to identify alternatively skipped exons (ASE), alternative 3′ splice sites (A3SS), alternative 5′ splice sites (A5SS), mutually exclusive exons (ME) and multiple exons skipping (Multi Skip) as previously described (*15*). A percent splicing index (PSI) value was calculated for each sample as the ratio of inclusion junction reads to the sum of inclusion and exclusion junction reads. As the datasets are paired, the difference in PSI values for each event (ΔPSI) was calculated as the median of ΔPSI values for each replicate. A filter is then applied on exon skipping events detected to select significant variants with an adjusted P value ≤0.05 and a ΔPSI value ≥10%. The gene expression level in each sample was calculated with HTSeq-count (v0.8.0) and differential gene expression between conditions was computed with DESeq2 R/ Bioconductor package (v1.10.1) (abs(log2FoldChange) ≥ 0.4 and 1.45, p ≤ 0.05) (*52*).

### Gene expression analysis by quantitative RT-PCR

Total RNA was extracted using the RNeasy Micro/Mini kit (Qiagen®) and reverse transcribed using random hexamers and Superscript III Reverse Transcriptase kit (Invitrogen®) according to the manufacturer’s protocol. Quantitative PCR reactions were carried out in 384-well plates using a QuantStudio 12K Flex Real-Time PCR System (Applied Biosystems®) with Power SYBR Green 2× Master Mix (Life Technologies®), 0.5 μl of cDNA, and 100 nmol/l of primers (Invitrogen®) in a final volume of 10 μl. Detailed information on the primers sequence is provided in Supplementary Table 3. Data were expressed as mean ± SD.

### Immunocytochemistry

After fixation with 4% paraformaldehyde (Euromedex®) for 15 minutes at room temperature, cells were incubated overnight at 4°C with the following primary antibodies : SQSTM1 (ab56416, ABCAM®), LAMP2 (ab25631, Calbiochem®), OCT3/4 (sc-5279 c-10, SantaCruz®), NANOG (D73G4, Cell Signalling®), SSEA3 (5600308, BD Pharmingen®), ISLET1 (AF1837, R&D Systems) and TUJ1 (PRB-435P, BioLegend). Appropriated secondary antibodies conjugated to Alexa fluorophores (Thermo Fisher Scientific®) together with Hoechst 33258 (5 μg/ml; Sigma-Aldrich®) were next applied to the cells. Images were acquired on a Zeiss inverted fluorescence microscope with the Zen blue software (Zeiss®) or with the HCS CellInsight CX7 device (Thermo Fisher®).

### Protein extraction and western blot analysis

Cells were lysed in RIPA buffer (Sigma-Aldrich®) containing 1% protease inhibitors (Sigma-Aldrich®) and 10% phosphatase inhibitors (Roche®). Proteins were quantified by BCA Protein Assay kit (Pierce®). Protein extracts (10 to 20µg) were loaded on 4–12% Nu-PAGE Bis-Tris gels (Invitrogen®) under reducing conditions and transferred to nitrocellulose membranes (Invitrogen®) using the iBlot2 Dry Blotting System (Invitrogen®). For LC3B protein analysis, PVDF Gel Transfer Stacks membranes were used (Invitrogen®). Membranes were next incubated with Blocking buffer (LI-COR®) supplemented with 0.1% Tween-20 and incubated overnight at 4°C with primary antibodies. Membranes were then incubated for 1 hour with the corresponding IRDye secondary antibodies (LI-COR®) and immunoreactive protein bands were detected using an Odissey CLx Imager (LI-COR®) according to the manufacturer’s protocol. Antibodies used in Western blotting analysis were anti-PLP1 antibody (ab28486, ABCAM®), anti-SQSTM1 antibody (ab56416, ABCAM®), anti-LC3B antibody (NB600-1384, Novus) and anti-ACTB antibody (92642210, LI-COR®).

### Prazosin treatment of zebrafish with sqstm1 knockdown

Zebrafish were microinjected with Morpholino antisense oligonucleotides (AMO) complementary to the zebrafish *SQSTM1* orthologue, sqstm1 (5′-ATGAAGAGACGGAAAGTGTCATCCT-3′) or with a control AMO (sqstm1-mis) (5′-ATCAACAGACCGAAACTCTCATCCT-3′), containing five mismatch nucleotides as previously described (*28*). 48 hpf embryos were submitted to Touch-evoked escape response test (TEER) prior to drug treatment and were after incubated overnight in 96-well plates. Prazosin was added to the embryo’s water at the final concentration of 10 μM. Spontaneous swimming between 6 and 48 h after the treatment was captured using a Zebralab system (ViewPoint, France) and analysed using the Fast Data Monitor software (ViewPoint, France). Embryos were recorded always at the same time and the experiment was repeated three different times. SQSTM1 expression level in zebrafish was analysis by electrophoresis and western blot using SQSTM1 (ab56416, ABCAM®) and ACTB (92642210, LI-COR®) antibodies.

### Statistical Analysis

Statistical analysis was performed using Student’s t-test to compare 2 groups. To compare 3 or more groups, we used parametric ANOVA or non-parametric (Kruskal-Wallis test) depending on the number of samples. When ANOVA or Kruskal-Wallis was significant, a post-hoc test was performed (see figure legends for details). A P value <0.05 was considered statistically significant. Analyses were performed using PRISM (version 5) software (GraphPad Software).

## List of Supplementary Materials

Fig S1 to S6.

Table S1 or Tables S3.

## Acknowledgments

We thank the FRANCE GENOMIQUE Platform for performing library preparation and sequencing. We gratefully acknowledge support from the PSMN (Pôle Scientifique de Modélisation Numérique) of the ENS de Lyon for the computing resources. The authors thank all the participants in the PULSE study and their families. The authors are grateful for financial support from the nonprofit research organization ARSLA (Christine Tabuenca, Marie France Cazalère, Marie Léon, Valérie Goutines and Sabine Turgeman), and for support from the French clinical research networks FILSLAN and ACT4ALS-MND, and the Fédération de la Recherche Clinique du CHU de Lille (Prof David Devos -coordinator of the Pulse study, Anne-Sophie Rolland, Alain Duhamel, Maeva Kheng, Julien Labreuch, Dominique Deplanque, Edouard Millois, Victor Laugeais, Maxime Caillier, Aymen Aouni, Pauline Guyon, Francine Niset, Valérie Santraine, Marie Pleuvret, Mathilde Bon and Laetitia Thibault) to the PULSE study. We thank Laetitia Barrault, Benjamin Brinon, Raphaël Woelke and Céline Buon (Team F Charbonnier, University Paris Cité & Inserm UMR_S1124, 75270, Paris Cedex 06, France) for their technical help. We express our gratitude to Pr Andrey Y. Abramov for providing SQSTM1 mutated fibroblasts. We express our gratitude to Pr Christine Baldeschi for providing adult normal keratinocytes. In memory of our friends and colleague Jacqueline Gide and Laetitia Barrault.

## Author contributions

Conceptualization: S.B., M.P. and C.M.; Methodology: S.B., M.C., A.B., C.B., E.K., M.P. and C.M.; Validation: S.B. and C.M.; Investigation: F.R., J.G, J.T., A.B., H.P., A.M., L.E.K., S.B; Resources: K.B., F.S., So.B., G.B., D.A., J.F.D., E.K., S.B., J.M.P.; Data Curation: H.P.; D.A.; Writing – Original Draft : S.B., M.P. and C.M.; Writing – Review & Editing Preparation: F.R., J.T., M.C., A.B., C.B., J.F.D., H.P., D.A., K.B., E.K., A.M., L.E., So.B., F.S., G.B., M.P., S.B. and C.M.; Supervision: M.P. and C.M.; Project Administration: M.P., S.B and C.M.; Funding Acquisition: C.M. and M.P.

## Data and materials availability

Data are available on GEO database (GSE261648).

---

**Suppl Fig. 1.**
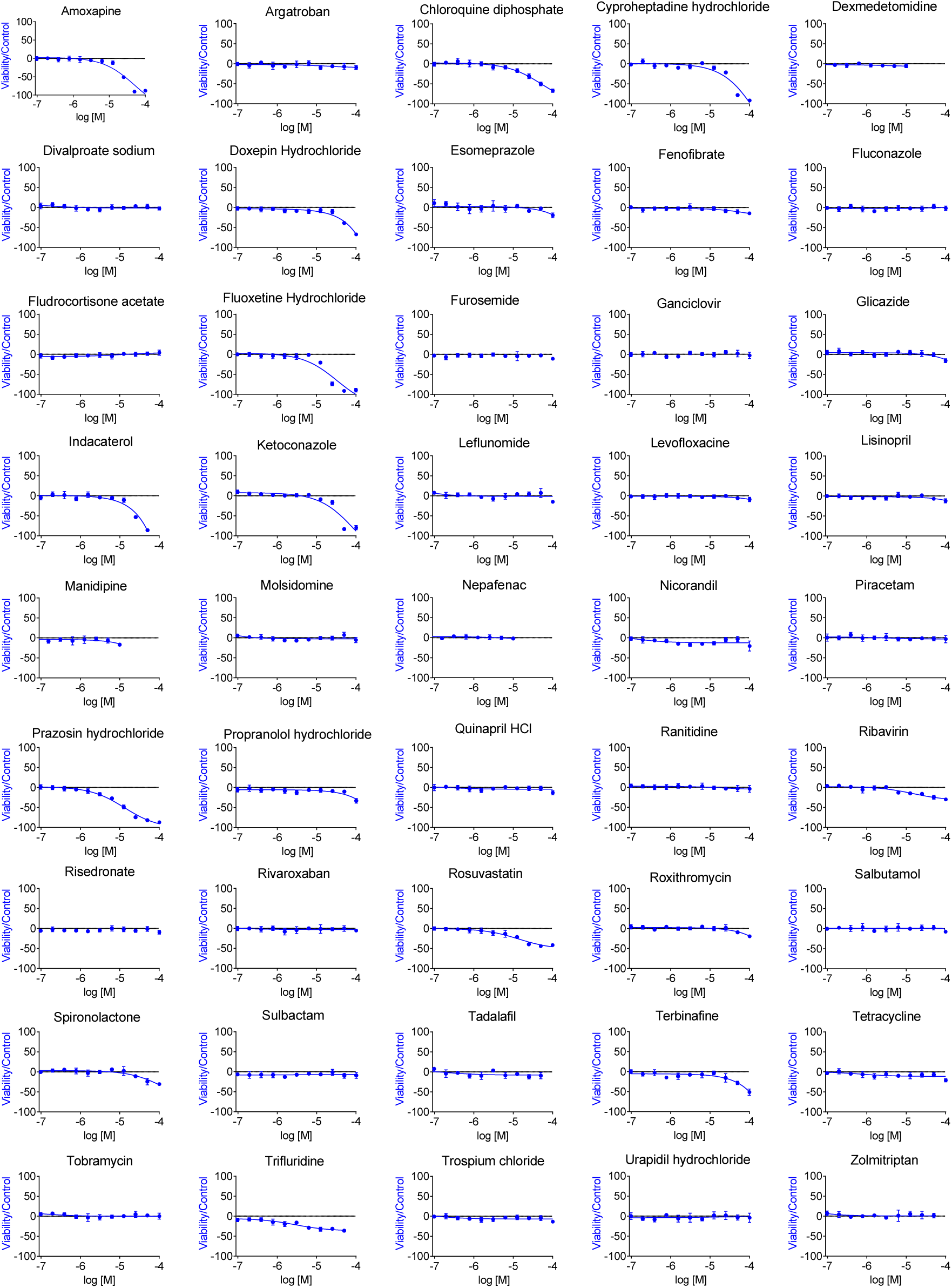
Viability of hES-derived MPCs treated with the compounds of the smart repositioning library. Cell viability of hES-derived MPCs was assessed using the Cell Titer-Glo assay following 24-hour treatment with various doses of drugs from the repositioning library.

**Suppl Fig. 2.**
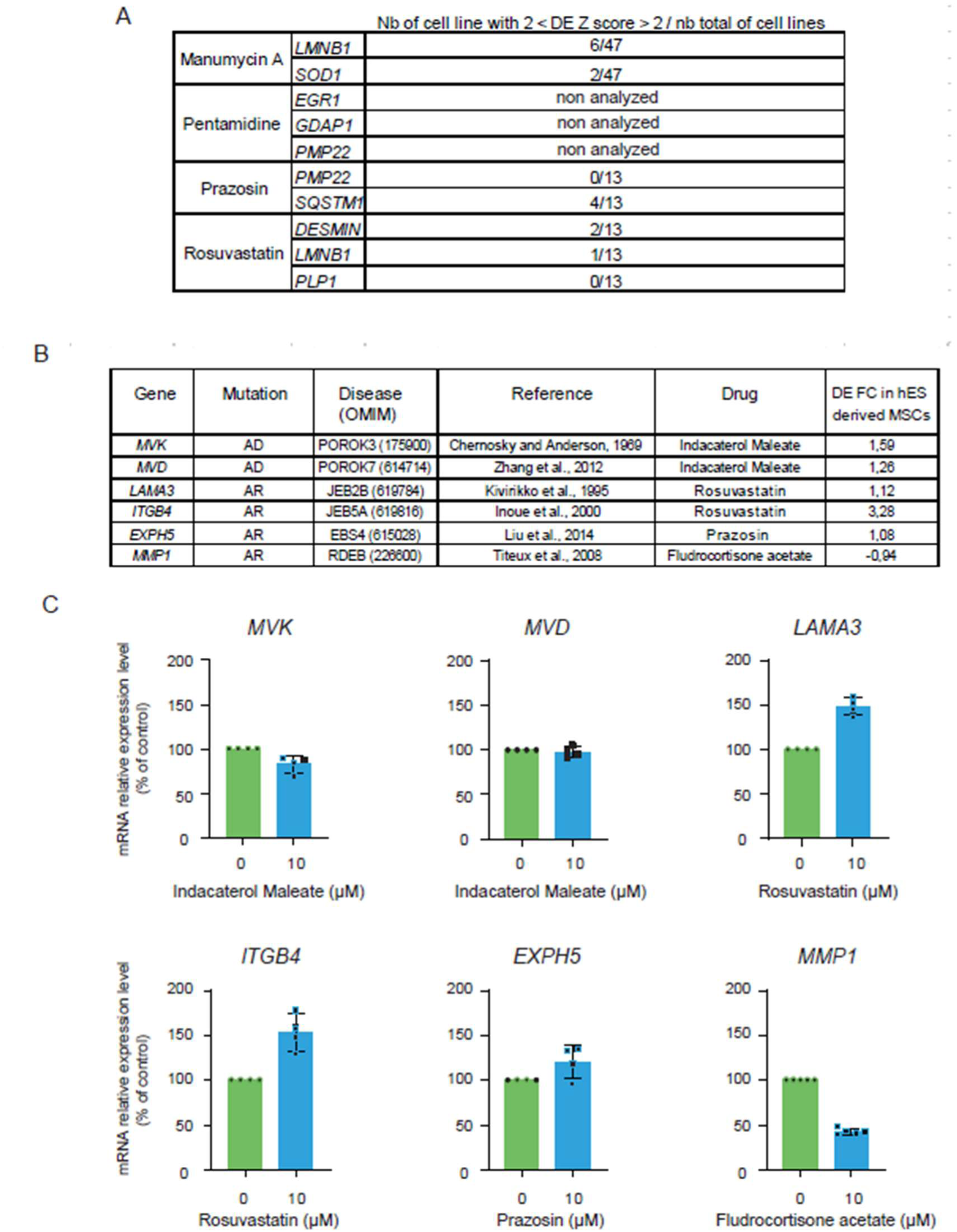
Analysis of gene expression regulation in various cellular contexts. **(A)** Gene regulations validated by RT-qPCR in hES-derived MPCs were analyzed using the Broad Institute’s Connectivity Map database. **(B)** Identification of six genes -*MVK* (mevalonate kinase), *MVD* (mevalonate diphosphate decarboxylase), *LAMA3* (laminin subunit alpha 3), *ITGBA* (integrin subunit beta 4), *EXPH5* (exophilin 5), and *MMP1* (matrix metallopeptidase 1) which are involved in genodermatoses and responsive to treatment with drugs from the smart drug library. **(C)** Plots of relative transcript expression for *MVK*, *MVD*, *LAMA3*, *ITGBA*, *EXPH5*, and *MMP1* in human adult keratinocytes, analyzed by RT-qPCR following 24-hour treatment with 10 µM of indacaterol maleate, rosuvastatin, prazosin, and fludrocortisone acetate. Data are presented as mean ± SD from three independent experiments (n = 3).

**Suppl Fig. 3.**
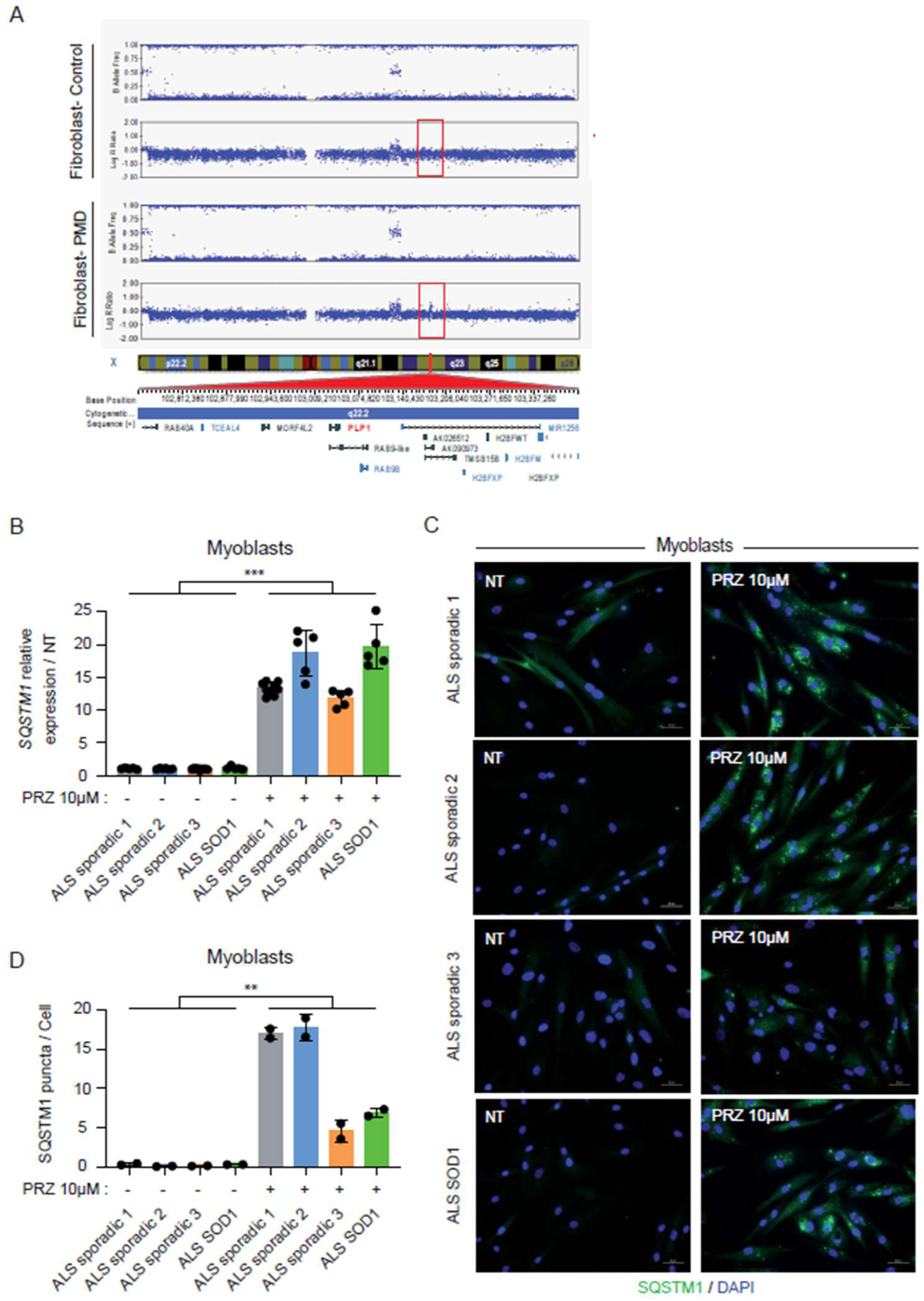
Characterization of PMD fibroblasts and ALS myoblasts. **(A)** SNP analysis of chromosome X in *PMD*-fibroblasts showing the amplification of a genomic region containing the *PLP1* gene (LOWER) compared to non-affected fibroblasts (UPPER). **(B)** Graphs of relative *SQSTM1* transcript expression in ALS-myoblasts, analyzed by RT-qPCR following 24-hour treatment with 10 µM prazosin. Data are presented as mean ± SD from three independent experiments (n = 3) and analyzed using Student’s t-test. **(C-D)** Representative immunocytochemistry images for DAPI (blue) and SQSTM1 staining (green) in ALS-mutated myoblasts 3021, 3055, 30620, and 3065. The number of nuclei and SQSTM1 puncta per cell were quantified after 24-hour treatment with 10 µM prazosin using automated microscopy acquisition and analysis. Data are shown as mean ± SEM from two independent experiments and analyzed with Student’s t-test (**P < 0.01, ***P < 0.001).

**Suppl Fig. 4.**
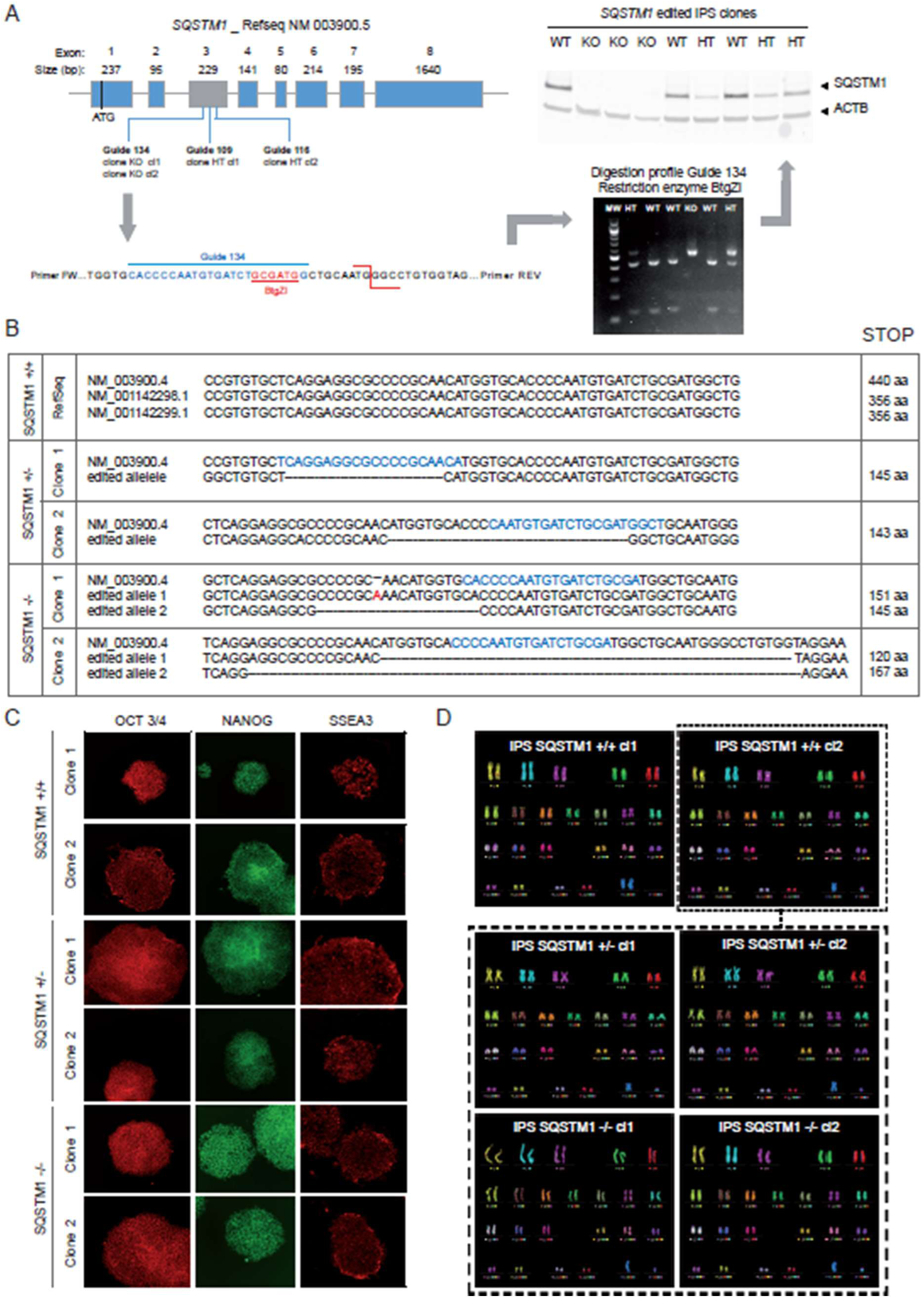
Generation of *SQSTM1* +/− and *SQSTM1* −/− hiPSC clones by the use of CRISPR/Cas9 technology. **(A)** Schematic representation of the editing strategy targeting Exon 2 of the *SQSTM1* gene. Guide 134 was used to generate two different *SQSTM1* −/− clones from 14c5 hiPSCs, while guides 109 and 116 were employed to obtain *SQSTM1* +/− hiPSC clones 1 and 2, respectively. Edited clones were selected based on their restriction profiles, confirmed by SQSTM1 Western blot analysis. **(B)** Sequences of *SQSTM1*-edited hiPSC clones show deletions or insertions on one or both alleles near the guide sequences (highlighted in blue). The position of the predicted premature STOP codon is indicated. **(C)** Representative immunocytochemistry images showing pluripotency markers OCT3/4 (red), NANOG (green), and SSEA3 (red) staining in control and edited 1869 hiPSC colonies. **(D)** Karyotype analysis of non-edited and edited *SQSTM1* hiPSC clones using mFish. *SQSTM1* +/− and *SQSTM1* −/− clones are derived from the hiPSC control clone 2 (cl2).

**Suppl Fig. 5.**
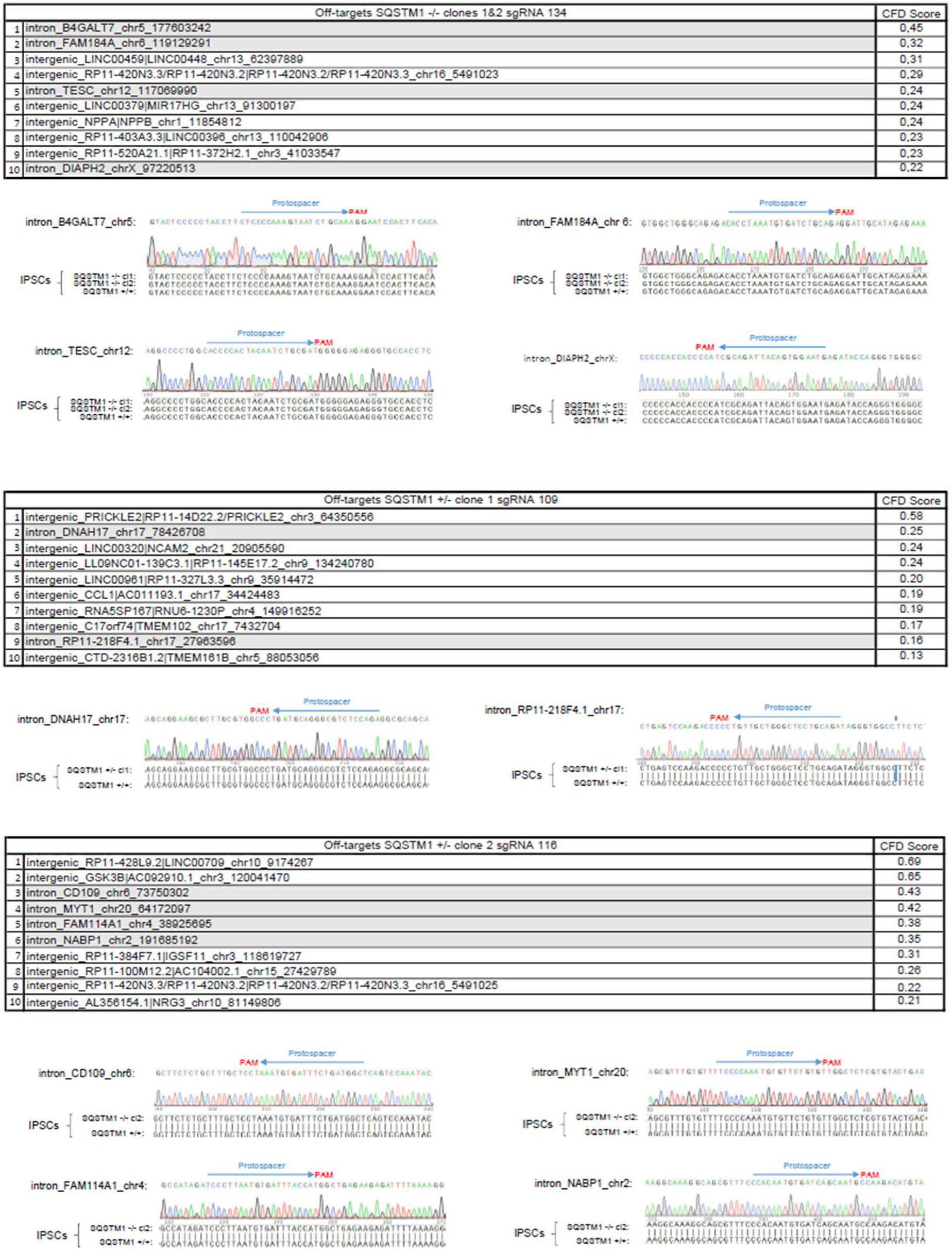
Off target analysis in edited *SQSTM1* +/− and *SQSTM1* −/− hiPSC lines. Potential off target edition sites in genomic DNA were identified with the CRISPOR (http://crispor.tefor.net/) software for the three guides used in the study. Region of interest located in within genes were amplified by PCR and analysed by Sanger sequencing. CFD means Cutting frequency determination indicated by the crispor software.

**Suppl Figure 6.**
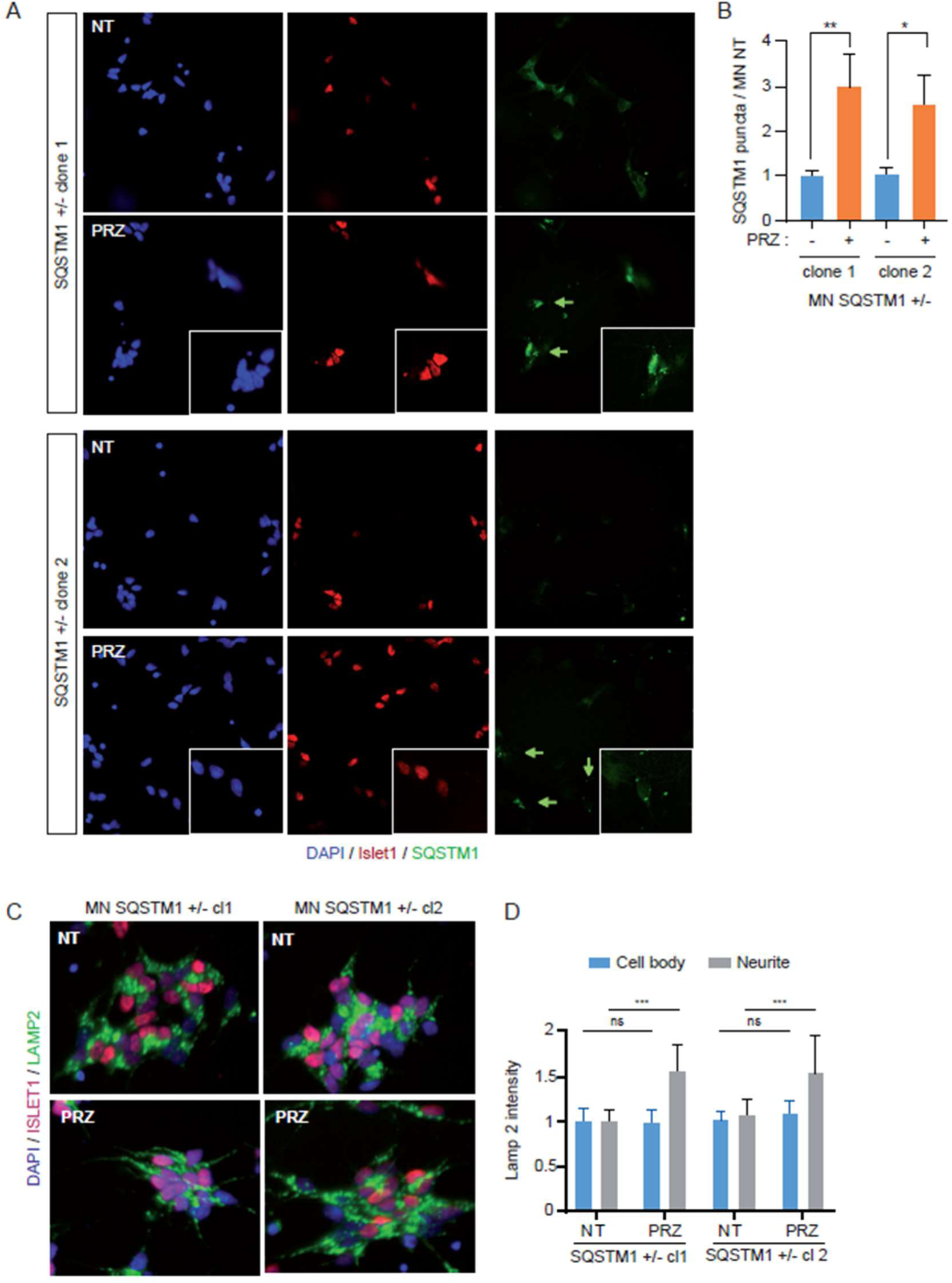
Effect of prazosin treatment in SQSTM1 +/− hiPSC-derived MNs on LAMP2 expression. (A-B) Representative images and quantification of immunocytochemistry for DAPI (blue), ISLET1 (red), and SQSTM1 (green) staining in motor neurons (MNs) at 24 days of differentiation, derived from two clones of SQSTM1+/− hiPSCs, either untreated (NT) or treated with 10 µM prazosin (PRZ) for 24 hours. SQSTM1 staining in SQSTM1+/− MNs treated with prazosin was normalized to the untreated condition. Data are presented as mean ± SD from three independent experiments (n = 3). Statistical significance was determined using the Kruskal-Wallis test with Dunnett’s post hoc multiple comparisons test (***P < 0.001). **(C-D)** Representative images of immunocytochemistry for DAPI (blue), ISLET1 (red), and LAMP2 (green) staining in SQSTM1+/− motor neurons treated for 24 hours with 10 µM prazosin. LAMP2 staining in the cell body or neurites of motor neurons was quantified by automated image acquisition and analysis. Data are shown as mean ± SD from three independent experiments (n = 3). Statistical analysis was performed using one-way ANOVA with Tukey’s post hoc multiple comparisons test (***P < 0.001).

**Supplemental Table 1.** List of the 50 small molecules tested for their repositioning potential.

| Drug | Synonyms | Activity | Famille/cible thérapeutique | Pubchem CID |
| --- | --- | --- | --- | --- |
| Cyproheptadine | Periactine | Anti-Allergic | Serotonin receptor antagonist | 2913 |
| Molsidomine | Corvasal | Anti-Anginal | Relaxation in the coronary blood vessels | 5353788 |
| Nicorandil | Ikorel | Anti-Anginal | Arterial and a venous dilator | 47528 |
| Indacaterol | Onbreezhaler | Anti-Asthmatic | Bronchodilator | 6918554 |
| Salbutamol | Ventoline | Anti-Asthmatic | Bronchodilator | 2083 |
| Levofloxacin | Tavanique | Anti-Bacterial | Fluoroquinolone antibiotic | 149096 |
| Manumycin | Manumycin A | Anti-Bacterial | Fatty amide | 6438330 |
| Roxithromycin | Rulid | Anti-Bacterial | Macrolide antibiotic | 6915744 |
| Sulbactam | Unasyn | Anti-Bacterial | Beta-lactamase inhibitor | 130313 |
| Tetracycline | Pylara | Anti-Bacterial | polyketide antibiotic | 54675776 |
| Tobramycin | Tobradex | Anti-Bacterial | Aminoglycoside antibiotic | 36294 |
| Argatroban | Arganova | Anti-Coagulant | Platelet Aggregation Inhibitors | 92722 |
| Rivaroxaban | Xarelto | Anti-Coagulant | Inhibitor of the coagulation factor Xa | 9875401 |
| Piracetam | Nootropyl | Anti-Convulsant | Neurotransmission promoter | 4843 |
| Amoxapine | Defanyl | Anti-Depressant | Norepinephrine and serotonin re-uptake inhibitor | 2170 |
| Doxepin | Quitaxon | Anti-Depressant | Norepinephrine and serotonin-reuptake inhibitor | 3158 |
| Fluoxetine | Prozac | Anti-Depressant | Selective serotonin-reuptake inhibitor | 3386 |
| Gliclazide | Diamicon | Anti-Diabetic | Insulin secretagogue | 3475 |
| Glyburide | Daonil | Anti-Diabetic | Insulin secretagogue | 3488 |
| Pioglitazone | Actos | Anti-Diabetic | Inducer of cellular responsiveness to insulin | 4829 |
| Divalproate sodium | Depakote | Anti-Epileptic | Gamma-aminobutyric acid (GABA) transaminase inhibitor | 18330669 |
| Fluconazole | Diffucan | Anti-Fungal | Triazole | 3365 |
| Ketoconazole | Ketoderm | Anti-Fungal | Phenylpiperazine | 456201 |
| Pentamidine | Pentacarinat | Anti-Fungal | Interfering with DNA replication | 4735 |
| Terbinafine | Lamisil | Anti-Fungal | Equalene epoxidase inhibitor | 1549008 |
| Furosemide | Lasix | Anti-Hypertensive | Potent loop diuretic | 3440 |
| Lisinopril | Zestril | Anti-Hypertensive | Angiotensin-converting enzyme (ACE) inhibitor | 5362119 |
| Manidipine | Iperden | Anti-Hypertensive | calcium channel blocker | 4008 |
| Prazosin | Minipress | Anti-Hypertensive | Alpha-1 adrenergic receptor inhibitor | 4893 |
| Propranolol hydrochloride | Avlocardyl | Anti-Hypertensive | Beta-adrenergic receptor blocker | 62882 |
| Quinapril HCl | Accupril | Anti-Hypertensive | Angiotensin converting enzyme (ACE) inhibitor | 54891 |
| Spironolactone | Aldactone | Anti-Hypertensive | Potassium-sparing diuretic | 5833 |
| Tadalafil | Cialis | Anti-Hypertensive | Cyclic guanosine monophosphate (cGMP)-specific type 5 phosphodiesterase inhibitor | 110635 |
| Urapidil | Eupressyl | Anti-Hypertensive | Adrenergic alpha-1 receptors inhibitor | 5639 |
| Risedronic acid | Risedronate | Anti-Hypocalcemic | Bisphosphonate | 5245 |
| Fludrocortisone acetate | Flucortac | Anti-inflammatory | Glucocorticoid-receptor agonist | 225609 |
| Indoprofen | Flosint | Anti-inflammatory | Prostaglandin-endoperoxide synthase inhibitor | 3718 |
| Leflunomide | Arava | Anti-inflammatory | Antirheumatical | 3899 |
| Nepafenac | Nevanac | Anti-inflammatory | Cyclooxygenase 1 and 2 inhibitor | 151075 |
| Fenofibrate | Lipanthyl | Anti-Lipidemic | Peroxisome proliferator activated receptor alpha (PPARalpha) activator | 3339 |
| Rosuvastatin | Crestor | Anti-Lipidemic | Hydroxymethyl-glutaryl coenzyme A (HMG-CoA) reductase inhibitor | 446157 |
| Chloroquine | Plaquenil | Anti-Malarial | Inhibit the parasitic enzyme heme polymerase | 2719 |
| Zolmitriptan | Zomig | Anti-Migraine | serotonin (5-HT) 1B receptors activator | 60857 |
| Esomeprazole | Innexium | Anti-Peptic | Gastric proton pump inhibitor | 9568614 |
| Ranitidine | Azantac | Anti-Peptic | Histamine H2-receptor antagonists | 3001055 |
| Tropium chloride | Ceris | Anti-Spasmotic | Muscarinic receptors blockade | 5284632 |
| Ganciclovir | Virgan | Anti-Viral | Analogue nucleoside | 135398740 |
| Ribavirin | Rebetol | Anti-Viral | Viral RNA synthesis inhibitor | 37542 |
| Trifluridine | Viropic | Anti-Viral | thymidylate synthase inhibitor | 6256 |
| Dexmedetomidine | Dexdor | Sedative | Alpha-adrenergic agonist | 5311068 |

**Supplemental Table 2.** Differentially expressed genes and splicing after drug treatment in MPCs.

| Molecule | DESeq log2FC $\geq 1,45$ | | | DESeq log2FC $\geq 0,4$ | | | Farline<br>Total<br>(somme) |
| --- | --- | --- | --- | --- | --- | --- | --- |
|  | DESeq up | DESeq down | Total | DESeq up | DESeq down | Total |  |
| DMSO | x | x | x | x | x | x | x |
| Amoxapine | 19 | 0 | 19 | 279 | 144 | 423 | 54 |
| Argatroban | 0 | 0 | 0 | 0 | 1 | 1 | 44 |
| Chloroquine | 31 | 1 | 32 | 758 | 383 | 1141 | 17 |
| Cyproheptadine | 11 | 0 | 11 | 312 | 89 | 401 | 54 |
| Dexmedetomidine | 0 | 0 | 0 | 0 | 0 | 0 | 27 |
| Divalproate sodium | 0 | 0 | 0 | 0 | 7 | 7 | 73 |
| Doxepin | 4 | 0 | 4 | 60 | 1 | 61 | 0 |
| Esomeprazole | 0 | 0 | 0 | 0 | 0 | 0 | 30 |
| Fenofibrate | 0 | 0 | 0 | 0 | 0 | 0 | 34 |
| Fluconazole | 0 | 0 | 0 | 0 | 0 | 0 | 31 |
| Fludrocortisone acetate | 3 | 0 | 3 | 108 | 162 | 270 | 31 |
| Fluoxetine | 17 | 0 | 17 | 239 | 93 | 332 | 32 |
| Furosemide | 0 | 0 | 0 | 0 | 0 | 0 | 47 |
| Ganciclovir | 0 | 0 | 0 | 0 | 0 | 0 | 15 |
| Gliclazide | 0 | 0 | 0 | 0 | 0 | 0 | 46 |
| Glyburide | 0 | 0 | 0 | 0 | 9 | 9 | 53 |
| Indacaterol | 24 | 0 | 24 | 385 | 132 | 517 | 47 |
| Indoprofen | 0 | 0 | 0 | 0 | 7 | 7 | 29 |
| Ketoconazole | 11 | 0 | 11 | 143 | 21 | 164 | 27 |
| Leflunomide | 0 | 0 | 0 | 25 | 38 | 63 | 21 |
| Levofloxacin | 0 | 0 | 0 | 0 | 0 | 0 | 23 |
| Lisinopril | 0 | 0 | 0 | 0 | 0 | 0 | 12 |
| Manidipine | 2 | 0 | 2 | 145 | 68 | 213 | 45 |
| Manumycin | 236 | 217 | 453 | 1508 | 1517 | 3025 | 125 |
| Molsidomine | 0 | 0 | 0 | 0 | 0 | 0 | 31 |
| Nepafenac | 0 | 0 | 0 | 0 | 3 | 3 | 36 |
| Nicorandil | 0 | 0 | 0 | 0 | 0 | 0 | 18 |
| Pentlseth | 124 | 128 | 252 | 1459 | 1446 | 2905 | 206 |
| Pioglitazone | 0 | 0 | 0 | 0 | 0 | 0 | 21 |
| Piracetam | 0 | 0 | 0 | 0 | 0 | 0 | 22 |
| Prazosin | 138 | 15 | 153 | 1418 | 1209 | 2627 | 70 |
| Propranolol hydrochloride | 0 | 0 | 0 | 0 | 0 | 0 | 27 |
| Quinapril HCl | 0 | 0 | 0 | 0 | 0 | 0 | 20 |
| Ranitidine | 0 | 0 | 0 | 0 | 0 | 0 | 10 |
| Ribavirin | 0 | 0 | 0 | 24 | 38 | 62 | 75 |
| Risedronic acid | 0 | 0 | 0 | 0 | 0 | 0 | 25 |
| Rivaroxaban | 0 | 0 | 0 | 7 | 11 | 18 | 48 |
| Rosuvastatin | 182 | 306 | 488 | 1724 | 1947 | 3671 | 140 |
| Roxithromycin | 0 | 0 | 0 | 17 | 3 | 20 | 17 |
| Salbutamol | 0 | 0 | 0 | 6 | 1 | 7 | 27 |
| Spironolactone | 0 | 0 | 0 | 46 | 41 | 87 | 51 |
| Sulbactam | 0 | 0 | 0 | 0 | 0 | 0 | 0 |
| Tadalafil | 0 | 0 | 0 | 1 | 0 | 1 | 44 |
| Terbinafine | 0 | 0 | 0 | 0 | 0 | 0 | 16 |
| Tetracycline | 0 | 0 | 0 | 0 | 0 | 0 | 47 |
| Tobramycin | 0 | 0 | 0 | 0 | 0 | 0 | 21 |
| Trifluridine | 3 | 0 | 3 | 348 | 59 | 407 | 57 |
| Tropium chloride | 0 | 0 | 0 | 0 | 0 | 0 | 34 |
| Urapidil | 0 | 0 | 0 | 0 | 0 | 0 | 49 |
| Zolmitriptan | 0 | 0 | 0 | 0 | 8 | 8 | 42 |

**Supplemental Table 3.**
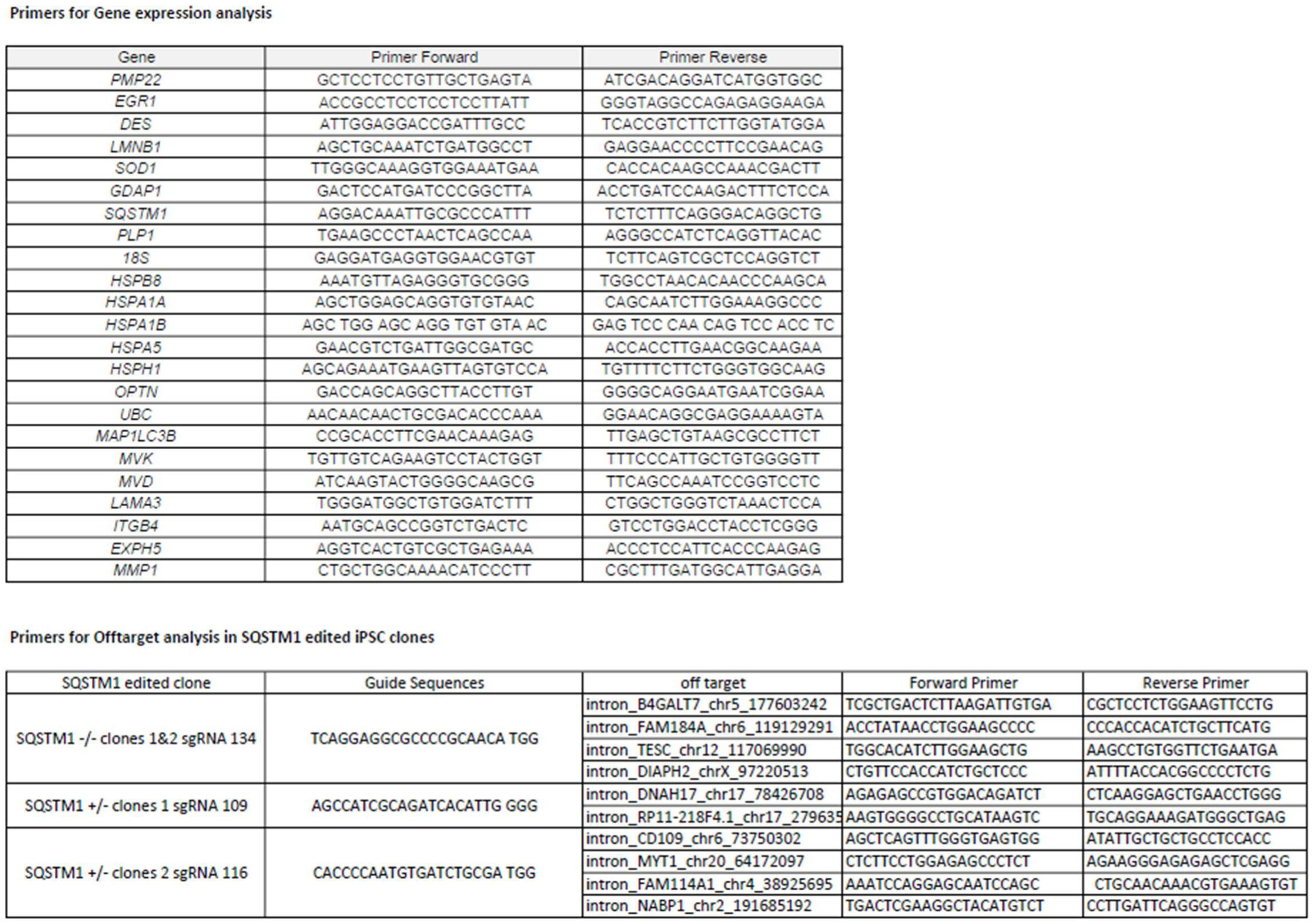
Primers for Gene expression and off target sequencing analysis.

